# Competition, precipitation and temperature shape deviations from scaling laws in the crown allometries of miombo woodlands

**DOI:** 10.1101/2024.10.30.621074

**Authors:** Arthur M. Yambayamba, Fabian J. Fischer, Tommaso Jucker

## Abstract

Scaling relationships between different axes of tree size, such as height, crown radius, crown depth and stem diameter, play a direct role in shaping forest structure and function. Theoretical models such as metabolic scaling theory postulate that they are optimized for biomechanical stability and hydraulic sap distribution. However, empirical data often show that such models are only good enough as first order approximations because they do not account for differences in species traits and environmental conditions where trees grow. Nevertheless, the vast majority of research has focused on temperate systems or tropical rainforests, so we continue to lack a full understanding of what factors shape allometries of trees in tropical dry forests. Here, we compile data on tree height, diameter, crown radius and depth from miombo woodlands across Zambia and use Bayesian hierarchical modelling framework to explore how allometric scaling relationships are shaped by climate and competition. Similar to previous studies, our results revealed that allometric scaling relationships deviate substantially from theoretical expectations. We found that competition, precipitation and temperature all affect crown allometric scaling relationships, with trees becoming more slender where neighbourhood competition was greater, while crowns were wider and deeper in warmer and wetter climates. Our study highlights how the structure and function of miombo woodlands is shaped by more than just water availability. Moreover, by developing improved crown allometric models for miombo woodlands, we provide new tools to aid the estimation of aboveground biomass and calibration of remote sensing products in these critically important dry forest ecosystems.

## Introduction

Allometric scaling relationships – which describe how an organism’s features scale with its size – are a cornerstone of ecology and have been used to capture everything from how organisms grow and move to the way energy flows through ecosystems (Huxley & Teissier, 1936; LaBarbera, 1989; Pélabon et al., 2014; Shingleton, 2010; Stillwell et al., 2016; West et al., 1997). In forest ecology, allometry has been used extensively to describe relationships between stem diameter at breast height (*D*), total tree height (*H*) and crown dimensions such as crown radius (*Crad*) and crown depth (*Cdep*), since these underpin our estimation of ecosystem functions and dynamics including carbon sequestration and storage (Coomes et al., 2012; Jucker et al., 2017; Lines et al., 2022; Shenkin et al., 2020). The first principles defining these relationships are founded on biomechanical, hydraulic and metabolic concepts that explore how *H*, *Crad* and *Cdep* scale with *D* (McMahon, 1973; Niklas & Spatz, 2004; Shinozaki et al., 1964; West et al., 1999). These scaling relationships assume that *D* must increase in proportion with *H* and the load it needs to support (i.e., crown size) in order to prevent a tree from buckling under its own weight or being blown over by wind (Eloy et al., 2017; King & Loucks, 1978; McMahon, 1973; McMahon & Kronauer, 1976). Similarly, assuming a fractal-like branching structure in trees, *D, H*, *Crad* and *Cdep* must scale with each other in such a way as to support photosynthetic organs, optimize hydraulic sap flow and minimize energy dissipation (Niklas & Spatz, 2004; Shinozaki et al., 1964; West et al., 1999; West et al., 2009).

Three theoretical models – metabolic scaling theory (MST), constant stress similarity (CSS) and geometric similarity (GS) – provide testable predictions on how *H*, *Crad* and *Cdep* should relate to *D* (Jucker et al., 2022; Niklas, 1995; Osunkoya et al., 2007; Shenkin et al., 2020; Sileshi et al., 2023). Specifically, these models assume that *D*, *H*, *Crad* and *Cdep* scale with each other following a power-law relationship with a fixed scaling exponent:

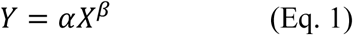

where *Y* is the dependent variable (e.g., *H, Crad, Cdep*), *X* is the independent variable (usually, *D* or *H*), α is the normalization constant (or log-scale intercept) and β is the scaling exponent (or log-scale slope). MST assumes that *H*, *Crad* and *Cdep* should scale with *D* in a way that not only ensures that a tree resists buckling under its own mass but also optimizes hydraulic sap distribution to the leaves. MST therefore postulates that *H, Crad* and *Cdep* should all scale to the 2/3 power of *D* (β = 0.67) while *Crad* should scale isometrically with *H* (β = 1) (Jucker et al., 2022; Shenkin et al., 2020; West et al., 1999; West et al., 2009). The CSS model instead predicts that *H* should scale to the 1/2 power of *D* (β = 0.50) so that trees do not exceed biomechanical constraints and avoid damage due to wind loads along their stem (Dean & Long, 1986; O’Brien et al., 1995; Osunkoya et al., 2007). Finally, the GS may be considered as a null model since the relative proportions of two related morphological variables (e.g. *D* and *H*) remains constant with tree size (Bertram, 1989; O’Brien et al., 1995). Therefore, the scaling exponent of *H*, *Crad* and *Cdep* with *D* and *Cdep* with *H* is predicted to β = 1.00 under the GS model.

However, several empirical studies have brought into question the generality of these theoretical models, especially when it comes to the predictions that MST makes about how *D*, *H*, *Crad* and *Cdep* should scale with each other (Coomes, 2006; Coomes & Allen, 2009; Harja et al., 2012; Muller-Landau et al., 2006; Russo et al., 2007). In particular, allometric scaling relationships have been observed to vary not just among species and biogeographic realms (Feldpausch et al., 2011; Jucker et al., 2022; Moncrieff et al., 2014; Shenkin et al., 2020), but also in relation to climate, soils, disturbance (e.g., wind, fire, herbivory, bark harvesting, logging) and local competition environment (Baldauf et al., 2021; Jackson et al., 2019; Jucker et al., 2014; Lines et al., 2012; Loubota Panzou et al., 2021; Moncrieff et al., 2011; Rutishauser et al., 2016). This suggests that while these theoretical models can serve as useful starting point from which to understand allometric scaling relationships in trees, universal scaling exponents have limited empirical support because trees are able to adjust the size and shape of their crowns depending on their micro-and macro-environment (Harja et al., 2012; Jucker et al., 2015; Lines et al., 2012).

Climatic factors such as solar radiation, precipitation and temperature directly influence tree growth and development. Tree allometries in different biomes therefore reflect their unique climatic conditions. It has been shown, for example, that trees become taller for a given *D* as water stress decreases (Chave et al., 2014; Feldpausch et al., 2011; Hulshof et al., 2015; Jucker et al., 2022). This suggests that hydraulic constraints are a major factor controlling *H*-*D* relationships in dry environments since taller trees are more vulnerable to hydraulic failure (Koch et al., 2004; Olson et al., 2018; Ryan et al., 2006; Ryan & Yoder, 1997; Stovall et al., 2019). By contrast, the effects of climate on allometric relationship between crown size (*Crad* and *Cdep*) and *D* appear much more variable among studies. For instance, a pantropical study on crown allometry revealed that for a given *D*, *Crad* and *Cdep* both decreased in wetter climates (Loubota Panzou et al., 2021). However, another pantropical study found no effect of precipitation on *Cdep*-*D* allometries (Shenkin et al., 2020). Another study across Spanish forests found that variation in precipitation and temperature had little or no effect on *Crad*-*D* scaling relationships within and across species (Lines et al., 2012). This suggests that other factors, including local competitive environment, are more important than climate in shaping crown allometry (Dieler & Pretzsch, 2013; Jucker et al., 2014; Jucker et al., 2015; Pretzsch, 2014; Seidel et al., 2011).

Competition for light, space, water and nutrients among neighboring trees all influence growth patterns and can therefore leave their fingerprint on crown allometry (McCarthy & Enquist, 2007). Generally, trees growing in dense, closed-canopy forests, where light is strongly limiting to growth, invest more in *H* for a given *D* to maximize light interception (Binkley et al., 2013; Harja et al., 2012; Lines et al., 2012). Conversely, trees in open-canopy forests are normally shorter for a given *D* than those in dense forests (Pommerening & Sánchez Meador, 2018). Furthermore, trees in open ecosystems normally have deeper *Cdep* for given *H* (Shenkin et al., 2020) and wider *Crad* for a given *D* (Loubota Panzou et al., 2021), as they are competing less intensely with neighbouring trees for space and light. However, when access to water becomes scarcer, trees may invest in smaller crowns to reduce transpiration and risk of hydraulic stress (Lines et al., 2012). Consequently, competition for light and water may modulate tree crown allometries in different ways depending on species traits, ontogeny and neighborhood environment (Coomes et al., 2011; Jucker et al., 2014; Niklas, 1995; Osunkoya et al., 2007). But their effects can be hard to disentangle because in drier environments where competition for water is stronger, forests also tend to be more open and less strongly influenced by competition for light.

Despite the plethora of studies that have characterized allometric scaling relationships in trees, the majority have focused on a single axis of crown allometry (most commonly *H*-*D* scaling relationships). Surprisingly, few studies have explored how multiple axes of crown allometry change in coordination with each other along climatic gradients and in response to competition with neighboring trees. And fewer still have done so in tropical dry forest ecosystems, despite the fact that these cover ∼0.5 billion ha of land (∼12% of global forest cover) and are unique in that they occupy climates that are seasonally dry and warm but still sustain relatively dense forest canopies (Ocón et al., 2021; Schröder et al., 2021).

Here we bring together data on *D*, *H*, *Crad* and *Cdep* for tens of thousands of trees distributed across Zambia’s miombo woodlands. Overall, these data represent 221 tree species and spans a broad range of climate (precipitation: 626-1678 mm yr^-1^ and temperature: 18.1-24.7°C) and competitive environments (stand basal area: 0.1-29.0 m^2^ ha^-1^) (Fig.1). Using these data, we sought to address three key questions relating to the allometry of trees in miombo woodlands:

**Fig. 1:**
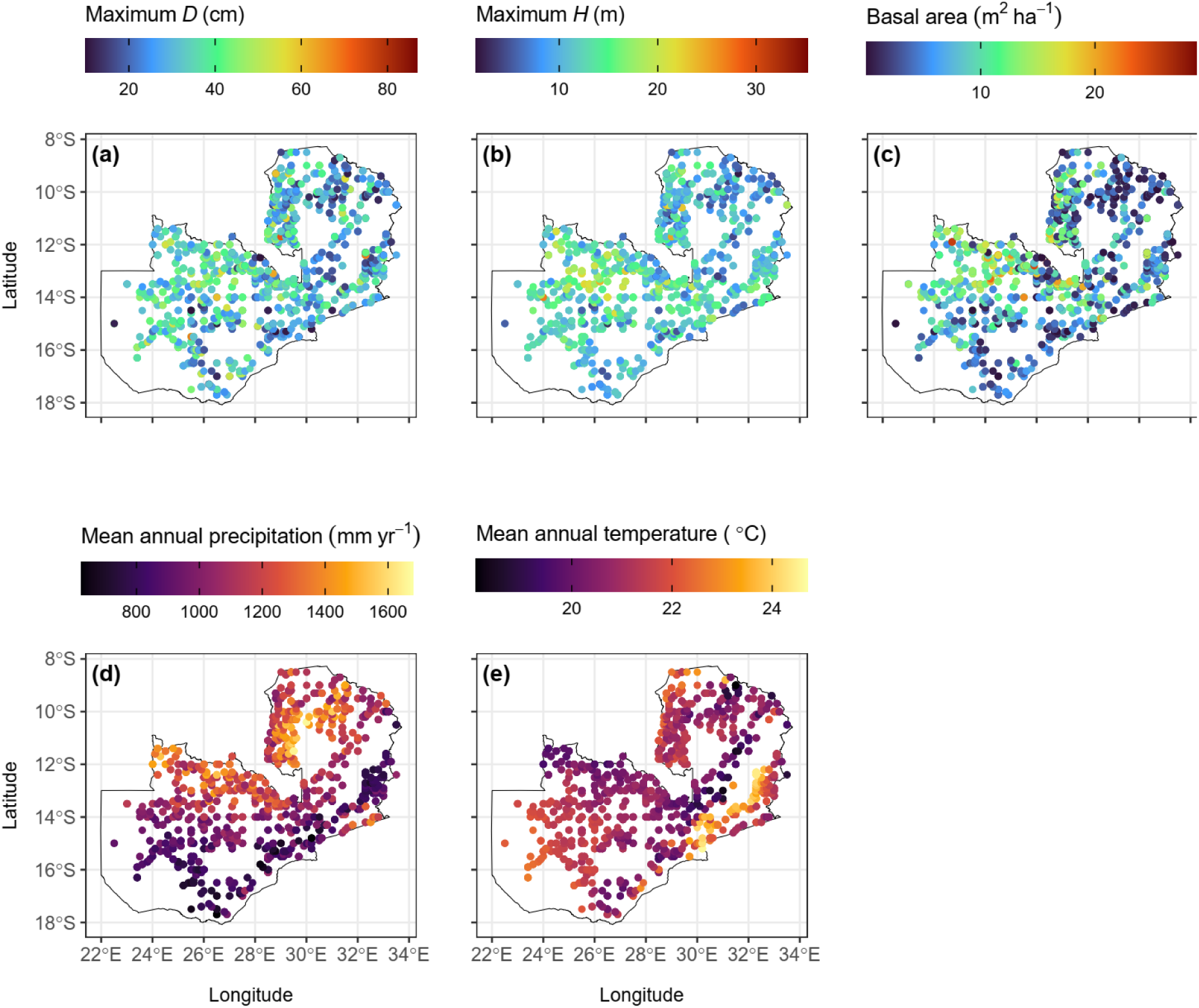
Variability in size structure and climate across the 1872 NFI plots in Zambia, including (**a**) maximum diameter at breast height (*D*), (**b**) maximum height (*H*), (**c**) basal area, (**d**) precipitation and (**e**) temperature. Climate data were extracted from the CHELSA-BIOCLIM+ database (Brun et al., 2022) at a spatial resolution of 30 arc-seconds (approximately 1 km).

(1.) How do theoretical allometric model predictions compare with observed scaling relationships between *H*, *Crad*, *Cdep* and *D*?

(2.) Do climate (temperature and precipitation) and competition (basal area) modulate community level allometric scaling relationships between *H*, *Crad*, *Cdep* and *D*?

(3.) How do *H*-*D* relationships vary among and within species along climate and competition gradients?

As byproduct of our analysis, we also intended to provide new and improved crown allometric scaling models for miombo woodlands, with potential applications for aboveground biomass estimation, the validation and calibration of rapidly growing remote sensing products of tree crown structure, and the development of better vegetation models (Chave et al., 2019; Fischer et al., 2019; Halperin et al., 2016; Jucker et al., 2017; Lines et al., 2022; Ngoma et al., 2019; Scheiter et al., 2013).

## Materials and methods

### Study area

Miombo woodlands are tropical dry forests that span large swaths of central and southern tropical Africa, covering an area of approximately 2 million km^2^ (Ribeiro et al., 2020). They are dominated by species in the Detarioideae subfamily of the Fabaceae, with the genera *Brachystegia* and *Julbernardia* being particularly prevalent (Chidumayo, 2013a; White, 1983), and mostly have either single-storied or two-storied stand structures (Chidumayo, 1987; Lawton, 1978).

Our study focused specifically on Zambia, where miombo woodlands cover >60% of the total forest area (ILUA, 2016). In terms of topography, Zambia is predominantly a plateau with miombo woodlands distributed approximately between 200–1800 m.a.s.l. in elevation. Zambia’s climate is characterized by three distinct seasons: hot-wet (mid-November to April), hot-dry (August to mid-November) and cool-dry (May to mid-August). Mean annual temperatures range between 19–25 °C while mean annual precipitation is around 600 mm in the south and 1400 mm in the northern part of the country. Based on the annual precipitation, the miombo woodlands can be divided into wet (>1,000 mm), intermediate (800-1,000 mm) and dry miombo (< 800 mm), which also correspond to the agro-ecological regions recognized in the country (Chidumayo, 2019; Chisanga et al., 2024; Chomba et al., 2012; Libanda et al., 2020). Soils are generally influenced by precipitation with more acidic and infertile soils in the wet miombo than in the intermediate-dry miombo, although the former has higher wood productivity (Chidumayo, 2019; ILUA, 2016).

Disturbances comprise land clearing for agriculture, wood harvesting for charcoal production and timber, wildfires and herbivory (mainly by elephants) (Chidumayo, 2013b; Ribeiro et al., 2020; Syampungani et al., 2016; Vinya et al., 2011). These activities have resulted in nearly 50% of protected forests in Zambia experiencing some form of disturbance in the last two decades (Phiri et al., 2023). This is consistent with a recent study on forest cover dynamics which revealed that primary forests decreased by 32% while secondary forests increased by 23% between 1972 and 2016 (Phiri et al., 2019) because most tree species in these woodlands have a high natural regeneration capacity and are able to recover after cessation of disturbances (Chidumayo, 2004; Syampungani et al., 2016). However, the effects of disturbance on forest stand structure and composition (e.g., tree density, basal area and species diversity) can persist for decades (Kalaba et al., 2013; McNicol et al., 2015; Poorter et al., 2021; Vásquez-Grandón et al., 2018).

### Field datasets

#### National Forest Inventory (NFI) dataset

For this study we compiled allometric data from two main sources (Table 1). The first were data from the National Forest Inventory (NFI) conducted between 2010–14 under the Integrated Land Use Assessment (ILUA) Phase II. NFI plots, each measuring 20×50 m (0.1 ha), were distributed across the whole of Zambia following a regular 1 km^2^ grid, with each grid cell containing a cluster of four plots. In each plot, all trees with *D* ≥ 10 cm were identified to species (or closest taxonomic unit) and their *D* and *H* were recorded using a diameter tape/caliper and *Suunto* hypsometers/range finder NIKON Forestry 550, respectively (Forestry Department, 2014; ILUA, 2016). Only plots where the vegetation type was recorded as miombo were retained for our analysis (Fig. S1). We excluded any trees where *D* < 9.5 cm (*n* = 366) and those where *D* > 100 cm, as there were only three of these with corresponding measurements of *H* and they represented a clear outlier in the size range of all other recorded trees. This left us with 32,804 unique tree records acquired across 1872 plots and belonging to 221 unique species and 124 genera (95.4% of trees identified to species; 1.1% to genus; 3.5% unidentified). The mean *D* of trees was 18.6 cm (range = 9.5–96.5 cm) and the mean *H* was 8.8 m (range = 1.3–37.0 m).

**Table 1:** Breakdown of the datasets used in the study, including number of plots, number of trees measured, number of species identified as well as the median and range (in square brackets) of diameter at breast height (*D*, in cm), total tree height (*H*, in m), crown depth (*Cdep*, in m), crown radius (*Crad*, in m), basal area (in m^2^ ha^-1^), tree/stem density, elevation (in m), mean annual precipitation (in mm yr^-1^) and mean annual temperature (in °C).

| Variable | NFI dataset | Crown size and shape dataset |
| --- | --- | --- |
| Number of plots laid out | 1,872 | 28 |
| Number of trees measured | 32,840 | 665 |
| Number of species identified | 221 | 4 |
| $D$ (in cm) | 15.8 [9.5 – 96.5] | 22.7 [10.4 – 64.0] |
| $H$ (in m) | 8.2 [1.3 – 37.0] | 15.9 [5.3 – 26.7] |
| $Cdep$ (in m) | - | 5.3 [1.10–12.5] |
| $Crad$ (in m) | - | 3.70 [1.15 – 11.33] |
| Basal area (in $\text{m}^2 \text{ha}^{-1}$) | 6.98 [0.08 – 29.02] | 12.00 [0.00 – 32.00] |
| Elevation (in m) | 1,199 [386 – 1,829] | 1,233 [1,007 – 1,313] |
| Tree density (stems $\text{ha}^{-1}$) | 210 [10 - 810] | - |
| Mean annual precipitation (in $\text{mm yr}^{-1}$) | 1,071 [626 – 1,678] | 1,051.5 [655 – 1,411] |
| Mean annual temperature (in $^{\circ}\text{C}$) | 21.0 [18.1 – 24.7] | 20.9 [20.0 – 22.0] |

#### Crown size and shape dataset

To complement the NFI dataset, which only contains information on *H* and *D*, in 2022–23 we established an additional 28 1-ha plots (100×100 m) where we also collected data on crown size and shape. These plots are distributed across wet (17 plots), intermediate (four plots) and dry miombo sites (seven plots) (Fig. S1) and also included primary (18 plots) and secondary (10 plots) forests. In each plot, we targeted at least 10 trees with *D* ≥ 10 cm for each of the dominant species (*Brachystegia boehmii*, *Brachystegia longifolia*, *Brachystegia spiciformis* and *Julbernardia paniculata*) in the stands. In plots with < 10 individuals per species, we sampled additional trees of the same species from the area just outside the plot.

For each tree, we measured *D* using a diameter tape, while *H* and *Cdep* (defined as total tree height – height to crown base) were measured using a Nikon Forestry Pro II Laser Rangefinder. We then measured *Crad* as the arithmetic mean of four crown radii measures in orthogonal cardinal directions using a measuring tape following RAINFOR protocols (Panzou & Feldpausch, 2020). In total, we measured 665 trees with a mean *D* of 25.2 cm (range = 10.4 – 64.0 cm), mean *H* of 15.6 m (range = 5.3–26.7 m), mean *Crad* of 4.0 m (range = 1.2 – 11.3 m), and mean *Cdep* of 5.5 m (range = 1.1–12.5 m).

### Disturbance and competition data

#### NFI dataset

For each NFI plot, we calculated several summary statistics relating to competitive environment and past disturbance history, including basal area (m^2^ ha^-1^), stem density (stems ha^-1^), maximum (95^th^ percentile) and median *D* and *H* (Poorter et al., 2021), but our main indicator of local competitive environment was basal area, as it incorporates both the effects of stem density and mean tree size (de Prado et al., 2022; Kuehne et al., 2019; Lines et al., 2012). In addition, we used the maximum and median *D* values to broadly classify each plot as either primary or secondary forest. Specifically, all plots with median *D* < 20 cm and maximum *D* < 30 cm were classified as secondary forests, while those where median *D* ≥ 20 cm and maximum *D* ≥ 30 cm were treated as primary forests. While only approximate, these thresholds were based on known size structure differences between primary and secondary miombo woodlands in Zambia (see Appendix S1 for details).

#### Crown size and shape dataset

For the 665 trees in the 28 1-ha plots in which we recorded the full suite of crown measurements, we estimated basal area surrounding each target tree using a Haglöf factor gauge with a basal area factor of 2 (Haglöf, 2014; Muir et al., 2011). Additionally, we classified each of the 28 plots into primary and secondary forests based on the same approach as for the NFI dataset.

### Climate data

Geographic coordinates were used to assign values of mean annual precipitation (mm yr^-1^) and temperature (°C) to all forest plots using the CHELSA-BIOCLIM+ database at a spatial resolution of 30 arc-seconds (approximately 1 km) (Brun et al., 2022). We chose mean annual precipitation and temperature as primary descriptors of climate because of their known influence on tree allometry and forest structure (Hulshof et al., 2015; Jucker et al., 2022; Lines et al., 2012). Moreover, as mentioned previously, mean annual precipitation is already widely used as a way to classify miombo woodlands into dry, intermediate and wet habitat types.

### Statistical analyses

We used a Bayesian hierarchical modelling approach to test the questions outlined in the introduction. All statistical analyses were done in R (version 4.3.1) using the brms, GGally, terra, tidyverse and data.table packages (Bürkner, 2017; Dowle & Srinivasan, 2023; R Core Team, 2023; Schloerke et al., 2021; Wickham et al., 2019).

#### Comparing empirical allometric exponents to theoretical predictions

To estimate the scaling exponents between different axes of crown allometry and compare these with theoretical predictions, we linearized the power law (Eq. 1) via a natural logarithm transformation as follows:

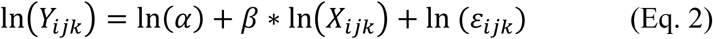

where *Y_ijk_* represents *H*, *Crad* or *Cdep* of tree *i* of species *j* in plot *k*, α is the normalization constant (log-scale intercept), β is the scaling exponent (log-scale slope), *X_ijk_* is either *D* or *H* of tree *i* of species *j* in plot *k* and ε_*ijk*_ is the random error associated with tree *i* of species *j* in plot *k* (see Table 2 for details).

**Table 2:** Estimates of the scaling exponents (β) for height-diameter-crown size allometries. The numbers in square brackets represent the 95% credible interval. *n*=number of trees; *H*=total tree height; *D*=tree diameter at breast height; *Crad* =crown radius and *Cdep*=crown depth.

| Allometric relationship | Estimate for the scaling exponent ($\beta$) using the NFI dataset | | | |
| --- | --- | --- | --- | --- |
| | All 221 species combined<br>($n = 32,804$) | *Brachystegia longifolia*<br>($n = 1,583$) | *Brachystegia spiciformis*<br>($n = 1,986$) | *Julbernardia paniculata*<br>($n = 2,816$) |
| *H-D* | 0.40 [0.39, 0.41] | 0.43 [0.41, 0.45] | 0.39 [0.37, 0.41] | 0.39 [0.37, 0.41] |
| Allometric relationship | Estimate for the scaling exponent ($\beta$) using the crown size and shape dataset | | | |
| | All 4 species combined<br>($n = 665$) | *Brachystegia longifolia*<br>($n = 174$) | *Brachystegia spiciformis*<br>($n = 139$) | *Julbernardia paniculata*<br>($n = 305$) |
| *H-D* | 0.37 [0.33, 0.41] | 0.49 [0.39, 0.58] | 0.35 [0.28, 0.41] | 0.35 [0.28, 0.41] |
| *Crad-D* | 0.81 [0.75, 0.87] | 0.66 [0.54, 0.78] | 0.89 [0.79, 1.00] | 0.73 [0.63, 0.83] |
| *Crad-H* | 0.60 [0.47, 0.72] | 0.55 [0.36, 0.74] | 0.83 [0.56, 1.11] | 0.50 [0.30, 0.70] |
| *Cdep-D* | 0.33 [0.25, 0.42] | 0.36 [0.19, 0.52] | 0.44 [0.31, 0.57] | 0.24 [0.11, 0.37] |
| *Cdep-H* | 0.78 [0.67, 0.90] | 0.59 [0.39, 0.79] | 0.76 [0.51, 1.00] | 0.75 [0.57, 0.92] |

i.) $H-D$ relationship for NFI data is based on the model: $\ln(H) \sim \ln(\alpha) + \beta * \ln(D)$ ; $\alpha \sim 1 + (1|\text{species}) + (1|\text{plot})$ ; $\beta \sim 1 + (1|\text{species}) +$ $(1|\text{plot})$ . ii.) The estimate for $\beta$ of the $H-D$ relationship (using the NFI dataset) for *Brachystegia boehmii* ( $n = 2,845$ ) was 0.40 [0.39, 0.42]. iii.) All the allometric relationships for the crown size and shape dataset are based on the model: $\ln(Y) \sim \ln(\alpha) + \beta * \ln(X)$ ; $\alpha \sim 1 + (1|\text{plot})$ ; $\beta \sim 1 + (1|\text{plot})$ ; where, $Y$ is $H$ or $Crad$ or $Cdep$ , $X$ is $D$ or $H$ , $\alpha$ is the normalization constant (intercept) and $\beta$ is the scaling exponent (slope).

Since we were interested in estimating scaling exponents for the entire dataset and for each species and plot, we modelled parameters for Eq. 2 as follows:

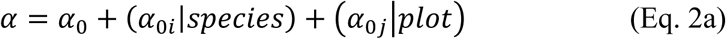

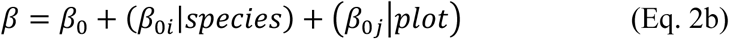

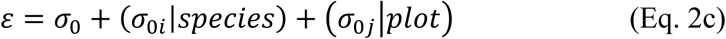

where α_0_, β_0_ and σ_0_ are population-level (fixed) effects; α_0*i*_, β_0*i*_ and σ_0*i*_ are species-level (random) effects; and α_0*j*_, β_0*j*_ and σ_0*j*_ are plot-level (random) effects for the normalization constant, scaling exponent and random error, respectively. Note that a small minority of trees in the NFI data were not identified to species level (4.6%) and were treated as one mixed-species group. For the crown size models, we did not include species as group-level (or random) effects since we only had four species as a priori fixed number for consideration. We therefore fitted one combined model with data from all species together, as well as separate models for the three species for which we had sufficient sample sizes (i.e., excluding *Brachystegia boehmii*, for which we only had data for 47 individual trees and because models for all allometric relationships converged with high numbers of divergent transitions).

*Testing the effects of climate and competition on tree crown allometric scaling relationships* To understand how climate and competition shape crown allometric scaling relationships in miombo woodlands, we expanded Eq. 2 (with only plot as a grouping factor) so that α and β were linear functions of basal area, precipitation and temperature:

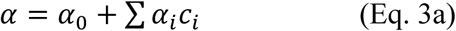

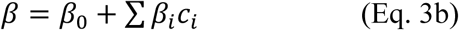

where α_0_ and β_0_are population-level (fixed) effects for α and β; α_*i*_ and β_*i*_are the coefficients for basal area, precipitation or temperature for α and β; and *c_i_* is basal area, precipitation or temperature, which were scaled and centered before fitting models so that their effect sizes could be directly compared. The estimated effect sizes therefore represented a change in α and/or β as a result of one-standard deviation increase in basal area, precipitation or temperature. Note that basal area, precipitation and temperature were only weakly correlated in our data (Fig. S5).

#### Effects of climate and competition on variation in H-D relationships within and among species

We used the NFI dataset to explore how *H-D* scaling relationships varied both within and among species depending on the climate and competitive environment. For this analysis we focused on four dominant tall-statured species (*Brachystegia boehmii*, *Brachystegia longifolia*, *Brachystegia spiciformis* and *Julbernardia paniculata*) and four dominant short-statured species (*Diplorhynchus condylocarpon*, *Pseudolachnostylis maprouneifolia*, *Uapaca kirkiana* and *Monotes africanus*) that are widely distributed across miombo woodlands (Chidumayo, 1987; Storrs, 1979; White, 1983) and together account for 42.4% of trees in the NFI dataset. Using these data we expanded Eq. 2 by allowing the normalization constant (α) and the scaling exponent (β) to vary within and across species as follows:

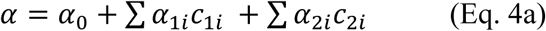

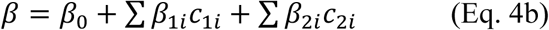

where α_0_ is the intercept for α and β_0_is the intercept for β; *c*_1*i*_is the species mean basal area, precipitation or temperature value (scaled and centered); α_1*i*_ and β_1*i*_ are the coefficients for the species mean basal area, precipitation or temperature for α and β respectively; *c*_2*i*_ represents the within species basal area, precipitation or temperature variability, measured as the deviation of each tree’s value from the group mean value, *c*_1*i*_ (Jucker et al., 2022); and α_2*i*_ and β_2*i*_ are the coefficients for the within species basal area, precipitation or temperature for α and β respectively.

#### Model fitting and parameter estimation

Parameter samples for all models described above were drawn using the no-U-turn sampler (NUTS), as implemented in the STAN software and exported to R via the brms package (Bürkner, 2017). Inference was based on 4000 posterior samples drawn from four parallel chains after discarding 1000 warmup draws per chain. For all model parameters, we used weak priors which we assumed to follow a normal distribution, with limits chosen so as to cover a realistic range of allometric parameters (Appendix 2).

Model convergence and fit was assessed via an estimate of the Gelman-Rubin convergence statistic (Rhat), the Bulk_ESS and Tail_ESS (effective sample size) statistics and posterior predictive checks. If a model has successfully converged, Rhat should be less than or equal to 1.01 while the Bulk_ESS and Tail_ESS should not be much lower (>10%) than the number of iterations (excluding warmups) (Bürkner, 2017; McElreath, 2020; Vanneste et al., 2024). All models successfully converged, with Rhat = 1.00 for each parameter, while both Bulk_ESS and Tail_ESS were >1000 in most cases, especially for population level estimates. Posterior predictive checks also revealed that the fitted models adequately reproduced the observed data (Figs. S6-S7).

## Results

### Testing the predictions of the GS, MST and STRESS models

#### Height-diameter scaling relationship

*H*-*D* scaling exponents were considerably lower than those predicted by MST and GS, and generally closer to the predictions of the CSS model (Fig. 2 and Table 2) across the entire NFI dataset with β = 0.40 [0.39, 0.41; 95% credible intervals] and similar estimates for the crown size and shape dataset (0.37 [0.33, 0.41]). We found that *H*-*D* scaling exponents varied among species, with *Brachystegia longifolia* having the highest exponent for both the NFI (0.43 [0.41, 0.45]) and crown size and shape datasets (0.49 [0.39, 0.58]), while *Diplorhynchus condylocarpon* had the lowest estimate (0.36 [0.33, 0.39]) among species recorded in the NFI dataset (Tables 2 and S1). We also observed considerable variation in scaling exponents among NFI plots, with values ranging from 0.48 [0.41, 0.55] to 0.28 [0.20, 0.37] (Table S2). However, we observed almost no systematic variation between either wet, intermediate and dry miombo woodlands, or between primary and secondary forests (Fig. 2). Similarly, there were no systematic patterns for different levels of disturbance or land cover classes (Figs. S8-S9).

**Fig. 2:**
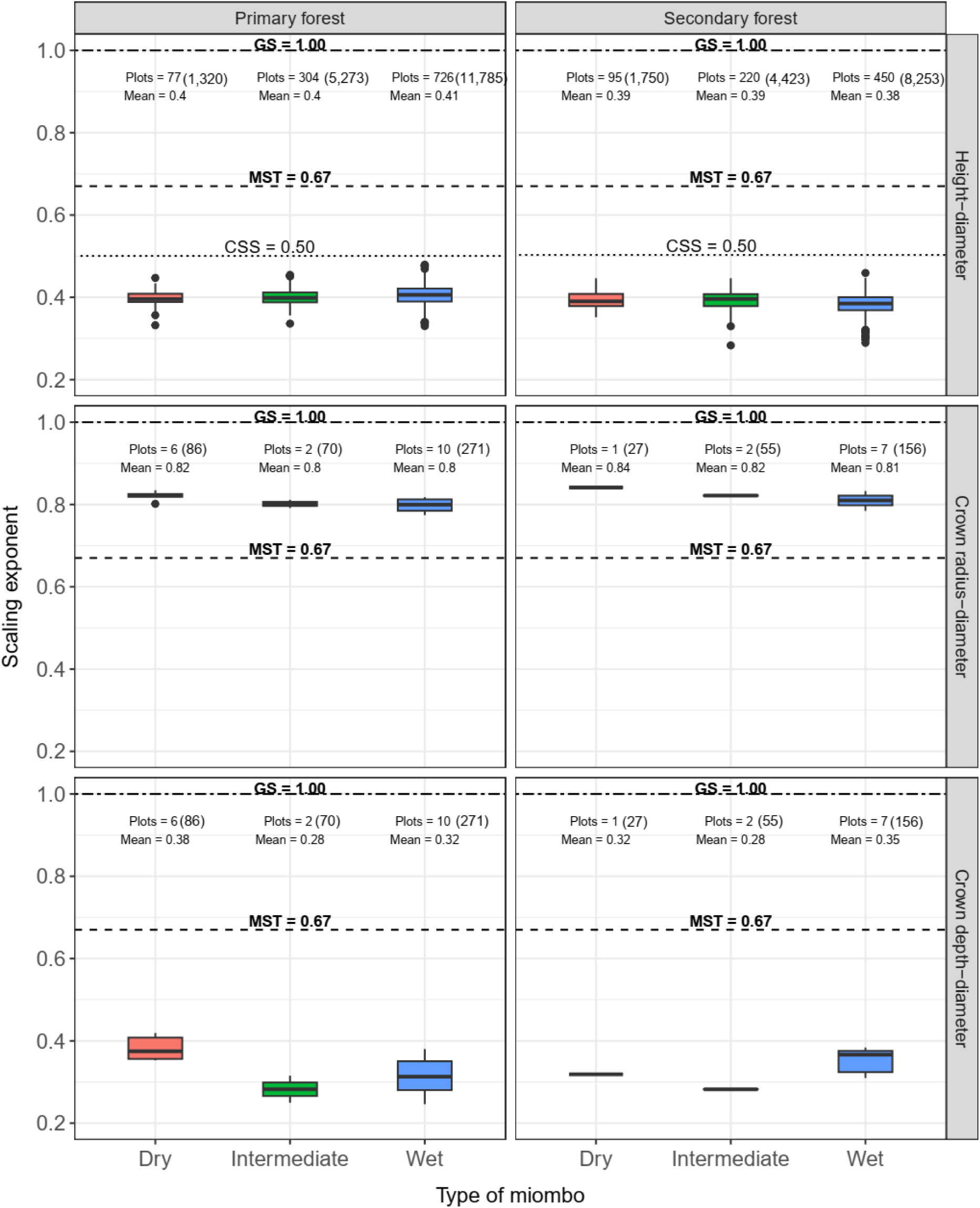
Variation in estimated scaling exponents for different allometric relationships in primary and secondary forests and across three types of miombo (dry, intermediate and wet). Theoretical predictions of MST (dashed; 0.67), CSS (dotted; 0.50) and GS (two dash; 1.00) are shown as horizontal lines in each panel. The number of trees in plots under different categories are shown in brackets beside number of plots.

#### Crown size scaling relationships

Overall, the *Crad*-*D* scaling exponent was close to, albeit higher (0.81 [0.75, 0.87]) than the MST prediction, and lower than that of GS (Fig. 2 and Table 2). However, species-level estimates of the *Crad*-*D* scaling exponent varied considerably, with both *Brachystegia longifolia* (0.66 [0.54, 0.78]) and *Julbernardia paniculata* (0.73 [0.63, 0.83]) overlapping with the predictions of MST, while *Brachystegia spiciformis* (0.89 [0.79, 1.00]) was not significantly different from the GS prediction (Table 2). We also found that *Crad*-*D* scaling exponents did not vary much in relation to forest or among miombo woodland types (Fig. 2 and Table S3b). When we expressed *Crad* as a function of *H*, we found that only *Brachystegia spiciformis* (0.83 [0.56, 1.11]) had an exponent that overlapped with the predictions of GS, while the other two had exponents that were all lower than MST predictions (Table 2).

As for crown depth, *Cdep*-*D* scaling exponents were all considerably smaller (0.33 [0.25, 0.42]) than MST and GS predictions (Fig. 2). However, *Cdep* scaling relationships were generally more variable than those for *Crad*, differing both among species and miombo woodland types (with higher scaling exponents in dry and wet forests). Similarly, *Cdep*-*H* scaling exponents (0.78 [0.67, 0.90]) where lower than the GS prediction. They also varied considerably among species (Table 2) and among miombo woodland types, with the highest exponents observed in forests classified as dry based on their precipitation (Fig. S10 and Table S3c).

### Influence of climate and competition on crown allometries

#### Height-diameter allometries

In the NFI dataset, we found that power-law scaling relationships between *H* and *D* were influenced mostly by basal area, and to a lesser degree by temperature and precipitation (Fig. 3). Shifts in the *H*-*D* scaling relationships were almost entirely due to changes in the normalization constant (α) term of the model, which increased with basal area (+0.05 [0.03, 0.07]), to a lesser degree, with temperature (+0.04 [0.03, 0.05]), and showed no effect for precipitation (0.00 [0.00, 0.00]). By contrast, the scaling exponent (β) was only affected by basal area (+0.02 [0.01, 0.03]) and largely unaffected by either precipitation or temperature (Fig. 3 and Table S4). As a result, a tree with *D* = 30 cm was on average 2.6 m (21.0 %) taller in a plot with high basal area compared to one with low basal area, 1.0 m (8.7 %) taller in warmer miombo woodlands compared to cooler ones, and 0.1 m (0.9%) taller in high precipitation area (Fig. 3g and Table S5a). These results were broadly consistent with those we obtained when modelling *H*-*D* scaling relationships with the much smaller crown size and shape dataset (665 trees from four species; Fig. S11). Here again we observed that trees were consistently taller in higher basal area plots, while temperature had a much smaller effect on height. However, in this second dataset we did observe a positive effect of precipitation on *H*-*D* scaling relationships, with the average tree with *D* = 30 cm being 1.4 m (8.1%) taller in plots with high compared to low precipitation (Table S5).

**Fig. 3:**
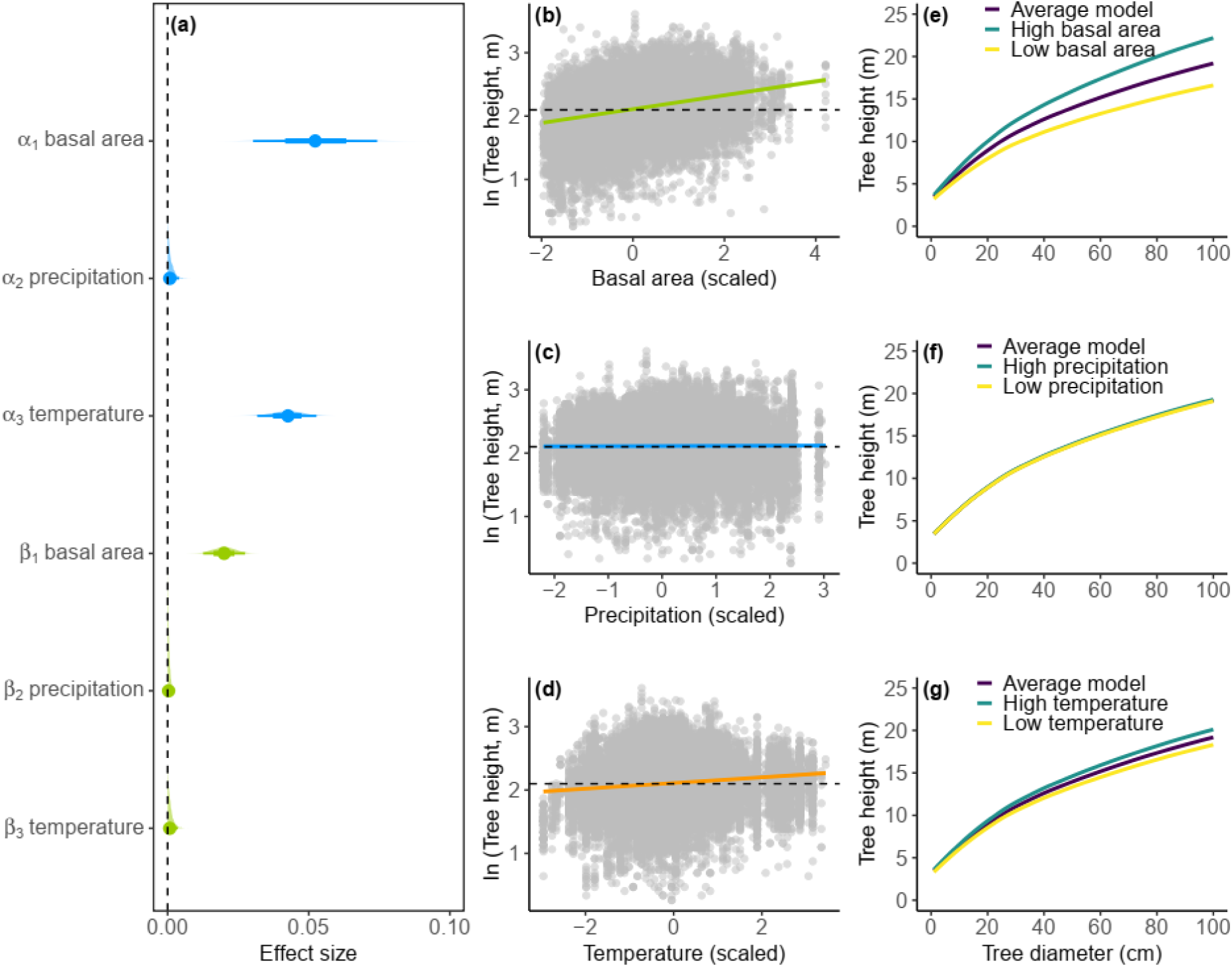
Effects of basal area, precipitation and temperature on **(a)** the height–diameter (*H-D*) power law model parameters, **(b**–**d)** *H* predictions for a tree of mean size along gradients of basal area, precipitation and temperature, and (**e**–**g**) *H* predictions across a range of *D* values (1-100 cm) while varying one model predictor (e.g., basal area in **e**) and keeping the other two (precipitation and temperature) constant at their mean value and setting their effects to zero. In (**b**–**d**), coloured lines represent mean model predictions (with shaded 95% credible intervals), the horizontal dashed line represents the mean tree *H* (on log scale) across the NFI dataset, and data points are shown as grey circles. In (**e**–**g**) low (yellow line) and high (blue line) prediction scenarios correspond to ± 1 standard deviation.

#### Crown size allometries

*Crad*-*D* scaling relationships were influenced more by basal area and temperature than precipitation, although in this case effect sizes were generally small (Fig. 4). Specifically, *Crad*-*D* power-law relationships became steeper as both basal area and temperature increased, with α decreasing (basal area = -0.15 [-0.30, -0.04]; temperature = -0.12 [-0.31, 0.03]) and β increasing (basal area = 0.04 [0.00, 0.08]; temperature = 0.05 [0.00, 0.11]); while precipitation effects were -0.05 [-0.16, 0.04] and 0.01 [0.00,0.04] for α and β respectively (Table S4). The average tree with *D* = 30 cm had a *Crad* that was 0.3 m (6.3%) wider in warmer miombo woodlands and 0.2 m (4.3%) narrower in more competitive environments but no change along the precipitation gradient (Fig. 4g and Table S6).

**Fig. 4:**
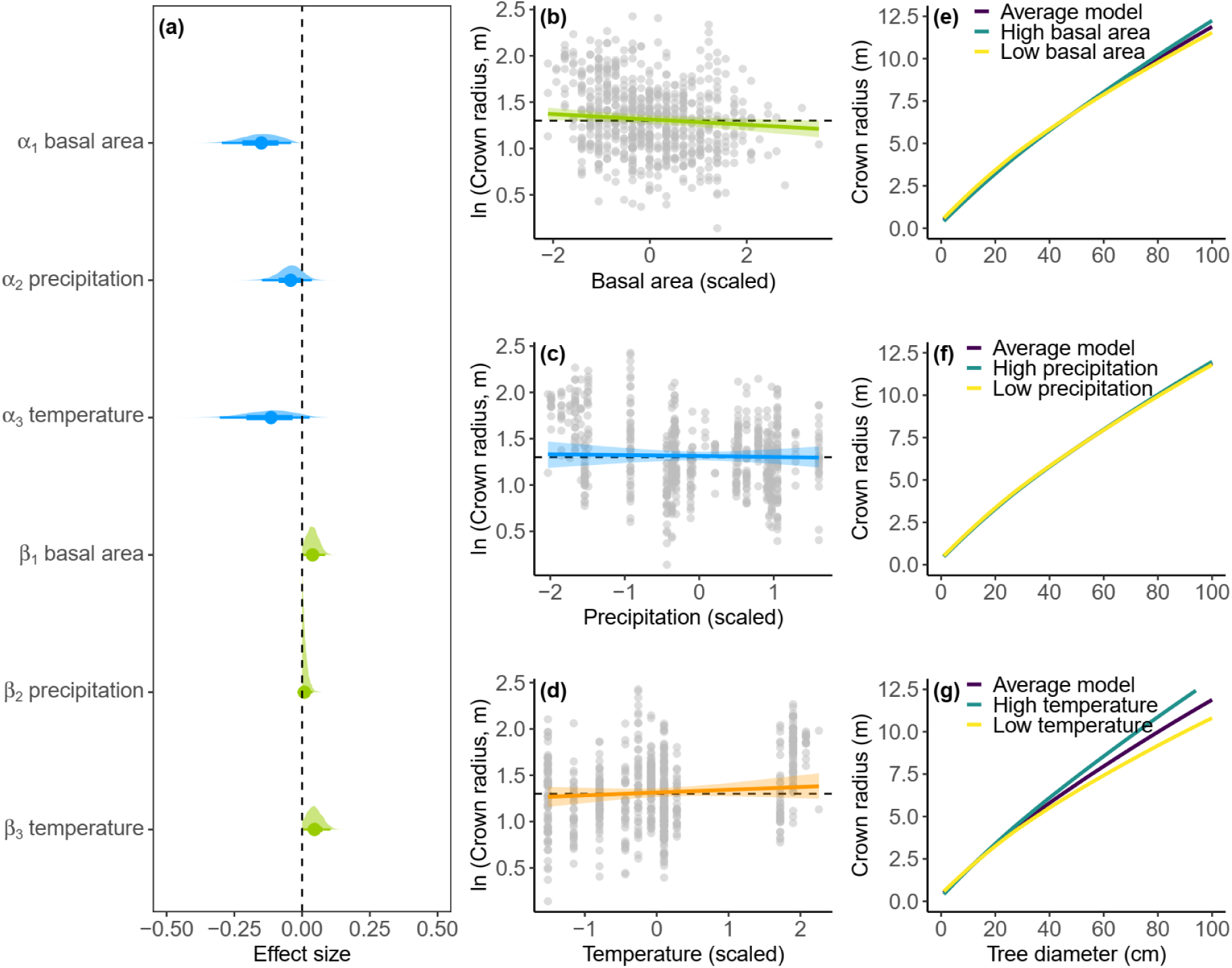
Effects of basal area, precipitation and temperature on **(a)** the crown radius–diameter (*Crad-D*) power law model parameters, **(b**–**d)** *Crad* predictions for a tree of mean size along gradients of basal area, precipitation and temperature, and (**e**–**g**) *Crad* predictions across a range of *D* values (1-100 cm) while varying one model predictor (e.g., basal area in **e**) and keeping the other two (precipitation and temperature) constant at their mean value and setting their effects to zero. In (**b**–**d**), coloured lines represent mean model predictions (with shaded 95% credible intervals), the horizontal dashed line represents the mean tree *Crad* (on log scale) across the crown size and shape dataset, and data points are shown as grey circles. In (**e**–**g**) low (yellow line) and high (blue line) prediction scenarios correspond to ± 1 standard deviation.

Instead, for crown depth we found that scaling relationships were much more strongly influenced by climate (Fig. 5). In particular, we found that *Cdep*-*D* power-law relationships became steeper as both temperature and precipitation increased, with α decreasing (temperature = -0.39 [-0.69, -0.10]; precipitation = -0.20 [-0.51, 0.06]) and β increasing (temperature = 0.18 [0.08, 0.27]; precipitation = 0.09 [0.01, 0.18]) (Table S4). As a result, the average tree with *D* = 30 cm had a crown that was 2.2 m (32.4%) deeper in warmer miombo woodlands (Fig. 5g and Table S7), and 1.1 m (17.7%) deeper at the wetter end of the distribution (Fig. 5f and Table S7). By contrast, basal area had a much smaller effect on crown depth, although we did find that crowns did became slightly deeper as basal area increased (Fig. 5e and Table S7).

**Fig. 5:**
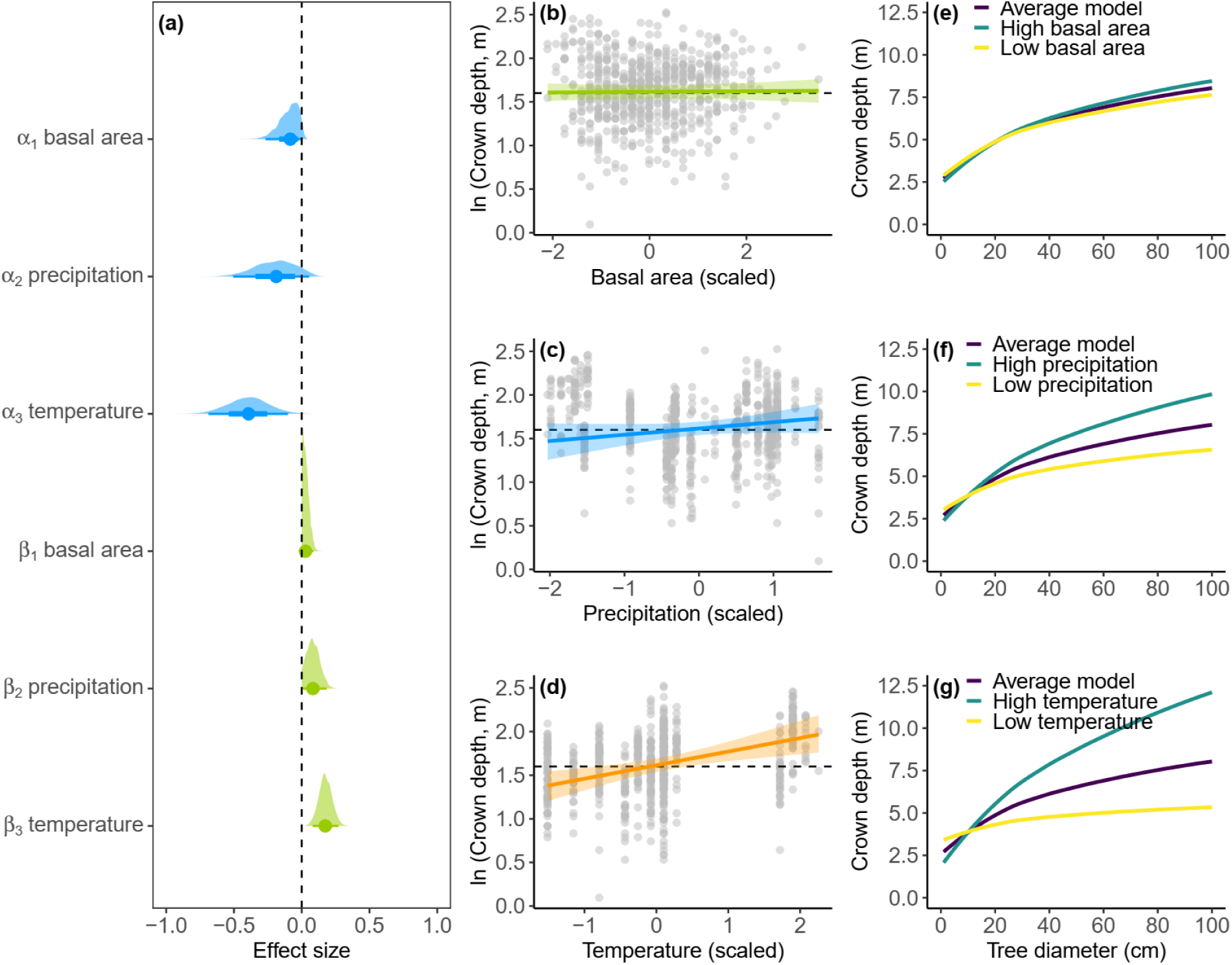
Effects of basal area, precipitation and temperature on **(a)** the crown depth–diameter (*Cdep-D*) power law model parameters, **(b**–**d)** *Cdep* predictions for a tree of mean size along gradients of basal area, precipitation and temperature, and (**e**–**g**) *Cdep* predictions across a range of *D* values (1-100 cm) while varying one model predictor (e.g., basal area in **e**) and keeping the other two (precipitation and temperature) constant at their mean value and setting their effects to zero. In (**b**–**d**), coloured lines represent mean model predictions (with shaded 95% credible intervals), the horizontal dashed line represents the mean tree *Cdep* (on log scale) across the crown size and shape dataset, and data points are shown as grey circles. In (**e**–**g**) low (yellow line) and high (blue line) prediction scenarios correspond to ± 1 standard deviation.

### Influence of climate and competition on inter- and intra-specific variability in height-diameter allometries among species

For the eight common species that make up > 40% of trees in the NFI data, we found similar shifts in *H*-*D* scaling relationships along gradients of basal area and temperature as those observed across species in the entire dataset. Specifically, all eight species became taller in plots with higher basal area (Fig. 6) and (to a lesser degree) where temperatures were higher (Fig. S15). By contrast, we found no effect of precipitation on variation in *H*-*D* scaling relationships within these common species, again confirming the general patterns that emerged across the entire dataset (Fig. S14). The effect of basal area on *H*-*D* scaling was consistent across the eight species, but appeared to be more pronounced in tall-stature species (average difference between low and high for the four species: + 2.9 m and + 21.9% in height for a tree with *D* = 30 cm; Fig. 6a-d) than short-stature ones (average difference between low and high for the four species: + 2.0 m and + 17.9% in height for a tree with *D* = 30 cm; Fig. 6e-h and Tables S9-S10). The effects of basal area on the power-law scaling relationship between *H* and *D*, both inter- and intra-specific, was almost exclusively related to increases in the normalization constant (Table S8 and Figs. S12-S13).

**Fig. 6:**
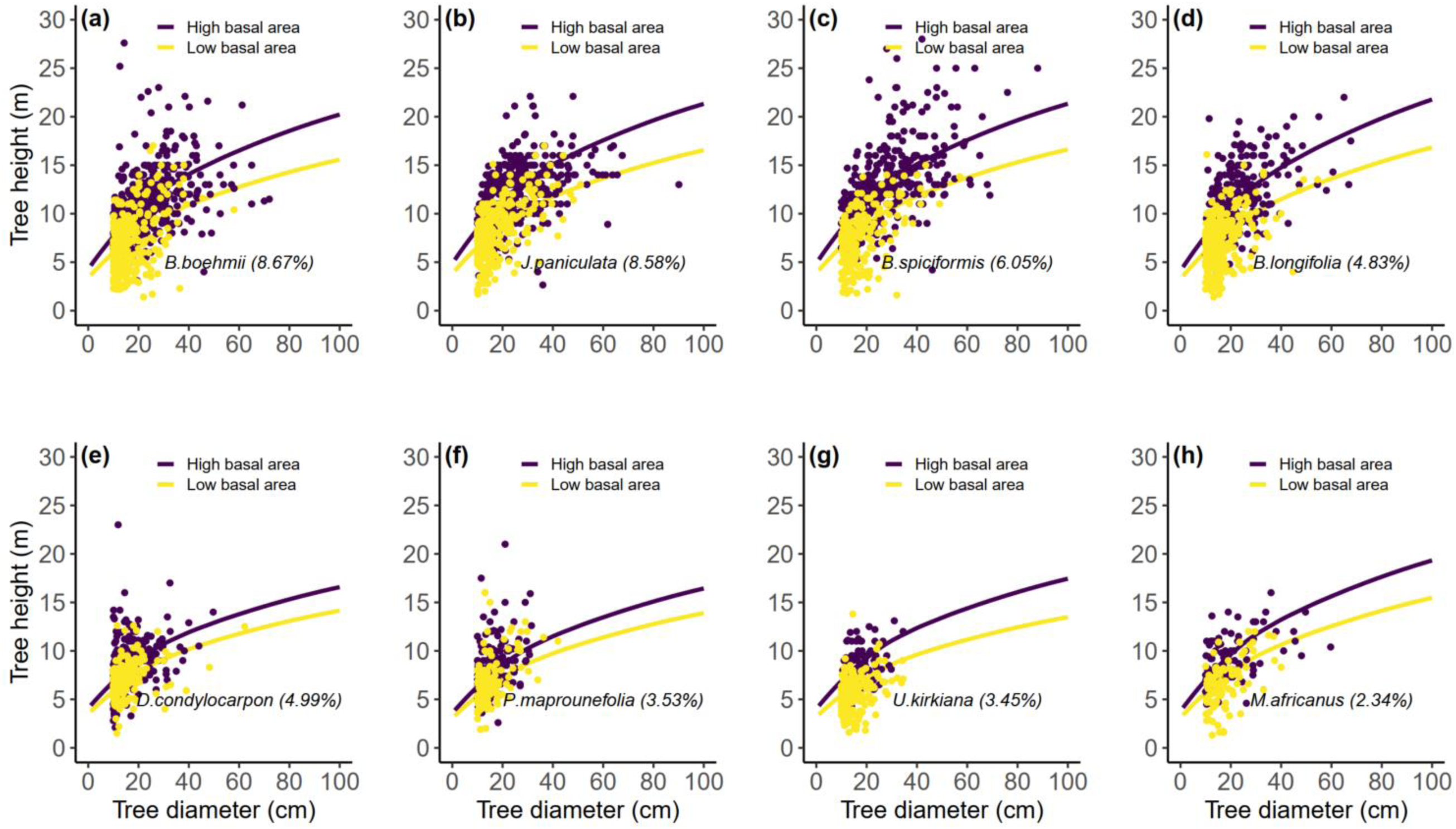
Effects of basal area on intra-specific variability in *H-D* scaling relationships for the four most abundant tall (**a**–**d**) and short (**e**–**h**) stature miombo tree species. Lines represent predictions for one-standard deviation increase (purple) and decrease (yellow) in basal area. Circles represent raw data from plots that are either one-standard deviation above (purple) or below (yellow) mean basal area. The percentage in brackets beside each species name corresponds to its relative abundance in the NFI data.

## Discussion

### Crown allometries in miombo woodlands deviate from theoretical expectations

*H*-*D* scaling in miombo woodlands, both across and within species, is closer to the CSS model prediction than the MST and GS. These results are similar to those of Jucker et al. (2022) for tropical dry forests and Shenkin et al. (2020) for pantropical forests, but in contrast to those of Handavu et al. (2021) who found good agreement with MST predictions for miombo woodlands on the Copperbelt Province of Zambia. Further, despite being very diverse in their growth strategies, the range for *H*-*D* scaling exponents is quite narrow among species. Nevertheless, tall-stature tree species had slightly higher scaling exponents than short-statured ones, with scaling exponents being positively correlated with tree height (Fig. S16). Our findings are also in line with hypotheses that allometric scaling exponents are constrained within narrow limits across evolutionary clades, taxonomic levels, geographical sampling areas, climate zones and disturbance regimes (Enquist & Niklas, 2001; Sileshi et al., 2023; Voje et al., 2014). We found nearly indistinguishable scaling exponents across the three types of miombo both within primary and secondary forests. Similarly, we did not observe any systematic variation in *H*-*D* scaling exponents with level of disturbance (Figs. 2, S8-S9). We thus argue that there is a tendency for *H*-*D* scaling exponents for different species within a given biome to cluster around some central value, even if this value does not conform to one of our current theoretical models.

Unlike for height, *Crad*-*D* scaling exponents are closer to the MST predictions. In fact, two of the three species-specific *Crad*-*D* scaling exponent estimates overlap with MST predictions. These findings are consistent with pantropical studies on crown size allometries (Blanchard et al., 2016; Loubota Panzou et al., 2021; Shenkin et al., 2020). However, *Crad*-*H* scaling exponents were well below the MST prediction of 1.00 – similar to findings by Jucker et al. (2022) – and tended to cluster around 0.67. Taken together, our data suggest that *Crad* in miombo woodlands (at least for the four dominant species studied here) scales with both *D* and *H* as 0.67. However, more data on crown radius from a range of species are needed to confirm this finding. By contrast, *Cdep* scaling relationships with both *D* and *H* showed more variation than *Crad*, and generally variation in scaling exponents among plots and species in *Cdep* was greater than any other scaling relationships considered (Table 2, Figs. 2 and S10). Our findings are similar to those of Shenkin et al. (2020), showing that *Cdep* scales with *H* with a power of 0.67, but not with *D* (Tables 2 and S3c). Further, the assertions that *H* (and subsequently *Cdep*, which is partially dependent on *H*) is more variable than *D* and *Crad* across ecosystems (Shenkin et al., 2020) is supported by our data.

### *H*-*D* scaling relationships are influenced more by competition than climate

We found that both climate and competition shape *H*-*D* allometric relationship in miombo woodlands, although the effect of the latter was significantly larger (Tables S4 and S8 and Fig. 3). The fact that trees grew taller for a given *D* is consistent with several other studies, including ones in seasonally dry Mediterranean forests (Antin et al., 2013; del Río et al., 2019; Feldpausch et al., 2011; Kafuti et al., 2022; Lines et al., 2012). Most miombo tree species, especially dominant ones, are light demanding (Syampungani et al., 2016) and hence taller in dense, undisturbed forests compared to more open, disturbed ones (Muvengwi et al., 2020) because of higher competition in the former. Trees growing in dense forests invest more in *H* growth than *D* during ontogeny in order to expose their crowns for better light capture (Antin et al., 2013). Our results also confirm that trees are shorter, for the same *D*, in open than closed canopy miombo woodlands (Munalula et al., 2020) because such environments offer less tree-tree competition for resources such as light and space. We argue therefore that human disturbances in miombo woodlands do not only alter the forest structure directly through the removal of trees, but also indirectly by altering local competitive environments. For example, it has been shown that selective logging leads to stands with shorter trees due to canopy gaps created which ultimately reduce competition for light (Rutishauser et al., 2016).

The weak effect of precipitation on *H-D* allometry in the NFI dataset is seemingly contrary to what we expected based on the literature and what we know about the role of precipitation in shaping the structure of miombo woodlands. Based on precipitation, miombo is divided into wet and dry woodlands and it is well established that trees are taller in the former (Frost, 1996; Munalula et al., 2020; White, 1983). However, precipitation did not have an effect on any of the *H*-*D* model parameters (when included in the model as the sole predictor variable or in combination with either basal area and/or temperature) (Figs. 3; S17-S20). There could be various explanations for our findings and here we present two suggestions. Firstly, while precipitation is responsible for high species turnover and productivity in the miombo (Chidumayo, 1987, 2019), competition, and not water availability, appears the stronger driver of the *H*-*D* scaling relationship. Therefore, for a given *D* and similar competitive environments, *H* would be indistinguishable between trees in wet and dry miombo. For instance, we observe nearly comparable forest structures (maximum *D*, *H* and basal area) between high and low precipitation in the northern and southern regions respectively (Fig. 1). The forest structure in the northern part may have been altered over the years due to fires and the *chitemene* (cut-and-burn) system of agriculture practiced in the area (Lawton, 1978; Stromgaard, 1986) leading to significant expansion of shorter secondary forests (Phiri et al., 2019). Our findings thus partly confirm those of Chidumayo (1987) showing that some traditionally known canopy and understory species can become canopy dominant in certain sub-regions of the miombo.

Another explanation for the seemingly unexpected results could be to do with inconsistencies in the underlying methodology used to measure *H*. Large-scale data collections such as the NFI program rely on a wide net of contributors who need to measure trees in difficult field conditions. While great care would have been taken to work with standardized protocols, their application in the field may have differed between groups. Given that more than 20 regional groups were involved in doing the measurements, there could be systematic under-or overestimation of tree *H* that mask the actual biological effects. Although this should in theory affect all the variables considered, precipitation effects would likely be the worst affected as rainfall varies considerably by sub-region so that teams that measured high precipitation areas would always have been different to those that measured low precipitation areas (Fig. 1). When we modelled *H-D* scaling relationships with our smaller crown size dataset (which was collected by the same field team), we did observe that trees became more slender in wetter areas, which is consistent with the literature (Jucker et al., 2022; Lines et al., 2012). However, even in this second dataset we still found that competition was a strong driver of *H-D* scaling relationships. Moreover, the crown size dataset was restricted to four tall-stature species and covered a much smaller gradient in precipitation than the NFI. Taken together, our findings suggest that while precipitation may play a key role in shaping *H-D* allometry in the miombo, we should consider the role of local competitive environment when modelling *H-D* scaling relationships in these woodlands.

In terms of temperature, we observed a marginal positive effect on *H* predictions for a given *D* in the NFI dataset. This was not entirely unexpected since the mean annual temperatures for miombo woodlands are well below estimated critical temperatures beyond which photosynthesis can be affected (Doughty et al., 2023; Winter, 2024). For tropical dry forests like the miombo, water availability, and not temperature, is the limiting factor for tree growth (Babst et al., 2019; Zuidema et al., 2022). In fact, using similar NFI dataset to ours, Godlee et al. (2024) found that higher temperatures during the rainy season in the miombo were associated with a longer growing season due to delayed leaf senescence. The marginal positive effect of temperature on *H* is thus plausible since some plots in wet miombo have higher mean annual temperatures than those in dry miombo (Figs.1 and S1, see also Munalula et al. (2020)). Conversely, higher temperatures coupled with low precipitation are associated with shorter trees for a given *D* (Chave et al., 2014).

### Crowns are wider and deeper in warmer, more open miombo woodlands

Our findings reveal that crowns became narrower for a given *D* in plots with higher basal area, while trees in warmer climates generally had wider crowns. By contrast, crown depth was little affected by competitive environment, and instead increased with both precipitation and temperature. Competition for light and space are known to drive crown allometries. The four species considered in our study are canopy-dominant species and therefore space could be one of the limiting factors for lateral crown expansion. It is thus not surprising that a combination of low competition (open canopy) and high temperature environments promoted larger *Crad* for a given *D*. This is also consistent with our results of slightly higher scaling exponents for both *Crad*-*D* and *Crad*-*H* (Figs. 2 and S10), and hence wider crowns, for trees in dry miombo (usually open canopy and higher temperatures) compared to wet miombo where competition is generally higher.

Our results thus provide empirical support to observations by Munalula et al. (2020) that trees in open canopy miombo are generally shorter and have wider crowns for a given *D*. This has also been demonstrated in pantropical studies comparing crown allometries in open and closed forest ecosystems (Loubota Panzou et al., 2021; Shenkin et al., 2020). Similar to our findings on *H*-*D* allometric relationship, temperature, and not precipitation, appears to have a stronger effect on both *Crad* and *Cdep* scaling relationships. We thus conclude that trees in warmer, open canopy miombo regions have wider and deeper crowns, while also being shorter – a crown architecture that would maximize light interception while minimizing hydraulic risk when competition for light is low.

## Conclusion

In this study, we have explored the relative effects of climate and competition in shaping tree allometries in miombo woodlands. While acknowledging the important role played by precipitation in tree growth, we find that competition and temperature exert larger effects on crown allometric scaling relationships. We therefore argue that allometric relationships in miombo should be viewed not only in the context of precipitation but also competition and temperature gradients. Consideration of these factors in sampling designs, development and application of allometric models should lead to more accurate estimations of *H* and crown dimensions. This should in turn lead to improved estimation of ecosystem structure, processes and functions, such as canopy cover, above ground biomass, forest productivity and tree volumes in miombo woodlands (Halperin et al., 2016; Handavu et al., 2021; Mauya et al., 2014; Ngoma et al., 2019). Further, field measurements of tree structure still remains a critical component for validation and calibration of emerging technologies such as LiDAR and related satellite biomass remote sensing missions (Chave et al., 2019; Lines et al., 2022). It is therefore imperative that we understand the drivers of landscape variations in tree structure such as demonstrated in our study.

## Acknowledgements

We are grateful for the funding made available through the Commonwealth Scholarship Commission to the first author. We acknowledge the Department of Forestry in the Ministry of Green Economy – Zambia for giving access to the NFI data. We are also grateful to the famers and institutions that gave access to their lands for collection of crown size and shape data. Further appreciation goes to the District Forestry Officers (in Mkushi, Kalulushi, Kapiri Mposhi and Kazungula Districts) who through their offices facilitated and/or helped with data collection. Last but not least, we thank all the field assistants, in particular Martin Mwiinga, who participated in data collection in all the plots. TJ was supported by a NERC Independent Research Fellowship (grant code: NE/S01537X/1). AMY was supported by PhD scholarship through the Commonwealth Scholarship Commission (CSC) programme (grant code: ZMCS-2021-542)

## Author contributions

AMY and TJ conceived the idea for the study. AMY did the data collection and led the analysis with assistance from FJF and TJ. All authors contributed substantially to revisions.

## Data availability statement

The NFI data has not yet been made publicly available and is therefore only available on request from the Department of Forestry of Zambia. Data on crown size and shape is available from the first author on request.

## Supporting information

Additional supporting information may be found online in the Supporting Information section.

