## Supplementary material for "Competition, precipitation and temperature shape deviations from scaling laws in the crown allometries of miombo woodlands": Yambayamba et al. 2024_supporting_information.docx

#### Appendix S1: Disturbance and competition data

For each NFI plot, we calculated a number of summary statistics relating to the competitive environment and past disturbance history, including basal area (m^2^ ha^-1^), stem density (stems ha^-1^), maximum (95^th^ percentile) and median $D$ and *H* (Poorter et al., 2021). We then used the maximum and median *D* estimates to classify each plot into either primary or secondary forest. All plots with median *D* < 20 cm and maximum *D* < 30 cm were classified as secondary and primary otherwise. This approach was adopted by inferring from literature and also arising from field experience by the first author. The minimum tree *D* allowed for harvesting as sawlog, and thus regarded as mature, in miombo is 30 cm. Furthermore, the mean *D* of trees in natural forests of Zambia is about 25 cm and thus the *D* class of 20–30 cm is regarded as medium (Kalinda et al., 2008). Secondary forests represent landscapes under transition through disturbance and regeneration (Phiri et al., 2019). Such forests are dominated by trees in *D* range of 5–20 cm (Chidumayo, 2019). We found that basal area was higher in primary than secondary forests. Further, dry miombo had the lowest basal area in both forest types (Fig. S2).

The NFI dataset included information on at most three environmental problems occurring in each plot and the level of intensity (classified as low, moderate, high and very high - with associated coding as 1, 2, 3 and 4 respectively) for each problem identified (Forestry Department, 2014). We extracted plots that had zero problems identified and those with one or two or three of the problems that included burning, overexploitation of forest resources and overgrazing since these are easier to observe and can be directly linked to human activities. This left us with 1170 (out of 1872) plots with record of discernible disturbance data. We then calculated the average intensity of disturbance of the identified problems and classified each plot accordingly as having zero, low, medium, high or very high intensity of disturbance. We found that basal area (m^2^ ha^-1^) was highest in plots with zero disturbance and lowest in plots classified as very high disturbance (Fig. S3). Furthermore, we found a consistent negative correlation between level of disturbance and basal area both across and within primary and secondary forests (Figs. S3 and S4a).

We also manipulated data on land cover classification for Zambia for the period of 2-years before and after the NFI was conducted. For this analysis, we used land cover maps for 2008 and 2016 to classify each plot into the nine land cover classes (primary forest, secondary forest, plantation forest, wetland, water body, cropland, irrigated crops, grassland, and settlement) based on the work of (Phiri et al., 2019). Using the two maps, we sought for consistent classification between the two time steps, i.e. plots being in same class for the two time steps. We found that no plot fell in the categories of plantation forest, water body and irrigated crops between the two time steps. Further, there were only two and five plots that fell in settlement and wetland categories respectively. We discarded these seven plots from the 1872 plots to make inference on disturbance and its relationship with basal area. We classified plots that did not maintain the same classification for the two time steps as transition. As expected, we found that basal area was highest in the primary forest and lowest in cropland land cover class (Fig. S4b).

#### Appendix S2: Statistical analyses

##### Motivation for using Bayesian regression methods

We used Bayesian hierarchical modelling approach to fit all the equations described in the main text. We chose Bayesian methods because they are robust in handling non-linear models and can facilitate simultaneous inter-specific and intra-specific inferences even for data-poor or under-represented species in the data (Cano et al., 2019; Tredennick et al., 2013). In the NFI dataset, we had 25 species, out of 221, with only a single tree recorded. Similarly, we had 66 plots with only a single tree recorded. This, however, did not pose challenges in models successfully converging when we specified plot and/or species as group-level (or random) effects.

The other reason we chose Bayesian methods is that the predictor variables we used are intuitively correlated; for instance, it is well established that increase in precipitation is accompanied by increased species and stem density and hence increased competition. Bayesian methods give robust estimates even in the presence of correlated variables since parameters are not estimated separately but rather from a joint distribution of parameters (Lines et al., 2012). We first scaled (centred) all the variables (basal area, elevation, precipitation, stem density (stems/ha) and temperature) we sought to consider as predictors and then ran a correlation test using the GGally R package version 2.1.2 (Schloerke et al., 2021; Zhou et al., 2023). We did not find strong correlation between climate and competition variables. However, precipitation and temperature were negatively correlated while basal area and stem density (stems/ha) were strongly positively correlated. Similarly, temperature and elevation were strongly negatively correlated (Fig. S5). We retained only variables that had less than 0.70 (absolute) correlation after running both the “pearson” and “spearman” cor methods in R (R Core Team, 2023). Hence, we excluded elevation and stems/ha from further analysis since these were strongly correlated with temperature and basal area respectively.

Lastly, Bayesian methods enabled us to generate posterior parameters distributions, for all plots and species in the NFI dataset, which can serve as baseline values for further works on description and modelling tree allometry in miombo woodlands.

##### Specification of priors

We assumed normal distribution for priors for all the parameters of the models fitted. We relied on literature values to set priors for $\alpha$ since the values for this parameter does not differ markedly for different studies. We used the MST predictions for $\beta$ to set the priors for the parameter. We used *N*(1.0, 1.0) for $\alpha$ and a truncated positive *N*(0.3, 0.5) for $\beta$. Priors for the species and plot levels were set in similar manner to correspond with parameters, i.e. $\alpha$ and $\beta$. We considered *N*(0, 0.5) for coefficients of basal area, precipitation and temperature and used *N*(0, 0.25) for the corresponding $\varepsilon$. Prior for the intercept of $\varepsilon$ was set at *N*(-1.0, 0.5).

##### Manipulation of expanded models and interpretation of results

We used estimated effect sizes for each variable (basal area, precipitation and temperature) to calculate $\alpha$ and $\beta$ (using equations 3 and 4) then ran predictions for *H*, *Crad* and *Cdep* for the *D* range of 1-100 cm at 1 cm intervals. We evaluated predictions in three different ways:

i.) Average model – where we used estimated model parameters ($\alpha_{0}$, $\beta_{0}$ and $\sigma_{0}$) of $\alpha$, $\beta$ and $\varepsilon$ with effect sizes for all other variables set to 0.
ii.) Low (basal area, precipitation or temperature) – where, for each variable separately, we calculated $\alpha, \beta\mathrm{and}\varepsilon$assuming a downward shift of one standard deviation compared to the average model (i.e. $\alpha_{0}$ - $\alpha_{i}$, $\beta_{0}$ - $\beta_{i}$ and $\sigma_{0}$ - $\sigma_{i}$). Coefficients of variables that were not analyzed were set to 0.
iii.) High (basal area, precipitation or temperature) – where, for each variable separately, we calculated $\alpha, \beta\mathrm{and}\varepsilon$assuming an upward shift of one standard deviation compared to the average model (i.e. $\alpha_{0}$ - $\alpha_{i}$, $\beta_{0}$ - $\beta_{i}$ and $\sigma_{0}$ - $\sigma_{i}$). Coefficients of variables that were not analyzed were set to 0.

Predicted *H*, *Crad* and *Cdep* values were back-transformed to the original scale (m) with the addition of the Baskerville correction factor ($\hat{\sigma}^{2}*0.5$)-where $\hat{\sigma}$ is the estimated residual standard error of the model (Baskerville, 1972).

### Supplementary tables

**Table S1:** Species-specific estimates of $\alpha$ and $\beta$ parameters for all the species in the NFI dataset. Microsoft Excell sheet

**Table S2:** Plot-specific estimates of $\alpha$ and $\beta$ parameters for all the plots recorded as miombo in the NFI dataset. Microsoft Excell sheet

**Table S3a:** Plot-specific estimates of $\alpha$ and $\beta$ parameters for all the plots in the crown size and shape dataset for *H-D* scaling relationship. Microsoft Excell sheet

**Table S3b:** Plot-specific estimates of $\alpha$ and $\beta$ parameters for all the plots in the crown size and shape dataset for *Crad-D* scaling relationship. Microsoft Excell sheet

**Table S3c:** Plot-specific estimates of $\alpha$ and $\beta$ parameters for all the plots in the crown size and shape dataset for *Cdep-H* scaling relationship. Microsoft Excell sheet

**Table S4:** Estimated mean [95% credible interval] of the model parameters, $\alpha$ and $\beta$ , and effect sizes of basal area, precipitation and temperature on $\alpha$ and $\beta$ of the power law model for the NFI dataset for *H*-*D* allometry and crown size and shape dataset for the *H-D*, *Crad*-*D* and *Cdep*-*D* allometries. Significant (excluding infinitesimal – where |0.00| is the lower and/or upper bound of the 95% credible interval) parameters and effect sizes are shown in bold. $\alpha$ estimates for all relationships and variables are on natural log scale. The effect size for each predictor variable represents the change in $\alpha$ and/or $\beta$ as a result of one-standard deviation increase in the predictor variable.

| Relationship | **Variable** |  | **Model parameter** | | |
| --- | --- | --- | --- | --- | --- |
|  |  |  | $\alpha$ Estimate [95% CI] |  | $\beta$ Estimate [95% CI] |
| *H*-*D*  (NFI dataset) | **Intercept** |  | **0.80 [0.78, 0.82]** |  | **0.46 [0.45, 0.47]** |
|  | **Basal area** |  | **0.05 [0.03, 0.07]** |  | **0.02 [0.01, 0.03]** |
|  | Precipitation |  | 0.00 [0.00, 0.00] |  | 0.00 [0.00, 0.00] |
|  | **Temperature** |  | **0.04 [0.03, 0.05]** |  | 0.00 [0.00, 0.00] |
| *H-D*  (crown size and shape dataset) | **Intercept** |  | **1.50 [1.37, 1.63]** |  | **0.38 [0.34, 0.43]** |
|  | Basal area |  | -0.01 [-0.08, 0.04] |  | 0.01 [0.00, 0.04] |
|  | Precipitation |  | 0.00 [-0.10, 0.08] |  | 0.01 [0.00, 0.04] |
|  | **Temperature** |  | **-0.13 [-0.26, -0.02]** |  | 0.03 [0.00, 0.08] |
| *Crad*-*D* | **Intercept** |  | **-1.16 [-1.36, -0.95]** |  | **0.78 [0.72, 0.85]** |
|  | **Basal area** |  | **-0.15 [-0.30, -0.04]** |  | 0.04 [0.00, 0.08] |
|  | Precipitation |  | -0.05 [-0.16, 0.04] |  | 0.01 [0.00, 0.04] |
|  | Temperature |  | -0.12 [-0.31, 0.03] |  | 0.05 [0.00, 0.11] |
| *Cdep*-*D* | **Intercept** |  | **0.67 [0.39, 0.94]** |  | **0.30 [0.21, 0.39]** |
|  | Basal area |  | -0.10 [-0.27, 0.00] |  | 0.03 [0.00, 0.09] |
|  | **Precipitation** |  | -0.20 [-0.51, 0.06] |  | **0.09 [0.01, 0.18]** |
|  | **Temperature** |  | **-0.39 [-0.69, -0.10]** |  | **0.18 [0.08, 0.27]** |

**Table S5a:** Predicted tree height, *H*, at different values of *D* using the fitted model for the NFI dataset and considering low and high basal area; low and high precipitation; and low and high temperature. Predictions of *H* in the table are based on equation 3 in the main text. Average model predictions are based on 0 effect sizes for all the predictors. Low and high represent one-standard deviation decrease (-1*mean effect size) and increase (+1*mean effect size) respectively, for basal area, precipitation and temperature.

| *D* (cm) |  | Predicted *H* (m) | | | | | | | | | |
| --- | --- | --- | --- | --- | --- | --- | --- | --- | --- | --- | --- |
|  |  | **Average model** |  | **Basal area** | |  | **Precipitation** | |  | **Temperature** | |
|  |  |  |  | Low | High |  | Low | High |  | Low | High |
| 10 |  | 6.6 |  | 6.0 | 7.3 |  | 6.6 | 6.7 |  | 6.3 | 6.9 |
| 15 |  | 8.0 |  | 7.2 | 8.9 |  | 8.0 | 8.0 |  | 7.6 | 8.4 |
| 20 |  | 9.1 |  | 8.2 | 10.2 |  | 9.1 | 9.2 |  | 8.7 | 9.6 |
| 25 |  | 10.1 |  | 9.0 | 11.4 |  | 10.1 | 10.2 |  | 9.7 | 10.6 |
| 30 |  | **11.0** |  | **9.8** | **12.4** |  | **11.0** | **11.1** |  | **10.5** | **11.5** |
| 40 |  | 12.6 |  | 11.1 | 14.3 |  | 12.5 | 12.6 |  | 12.0 | 13.2 |
| 50 |  | 13.9 |  | 12.2 | 15.9 |  | 13.9 | 14.0 |  | 13.3 | 14.6 |
| 60 |  | 15.2 |  | 13.3 | 17.4 |  | 15.1 | 15.2 |  | 14.5 | 15.9 |
| 70 |  | 16.3 |  | 14.2 | 18.7 |  | 16.2 | 16.4 |  | 15.6 | 17.1 |
| 80 |  | 17.3 |  | 15.1 | 20.0 |  | 17.2 | 17.4 |  | 16.5 | 18.2 |
| 90 |  | 18.3 |  | 15.9 | 21.1 |  | 18.2 | 18.4 |  | 17.5 | 19.2 |
| 100 |  | 16.1 |  | 14.1 | 18.1 |  | 16.0 | 16.2 |  | 15.4 | 16.8 |

**Table S5b:** Predicted tree height, *H*, at different values of *D* using the fitted model for the crown size and shape dataset and considering low and high basal area; low and high precipitation; and low and high temperature. Predictions of *H* in the table are based on equation 3 in the main text. Average model predictions are based on 0 effect sizes for all the predictors. Low and high represent one-standard deviation decrease (-1*mean effect size) and increase (+1*mean effect size) respectively, for basal area, precipitation and temperature.

| *D* (cm) |  | Predicted *H* (m) | | | | | | | | | |
| --- | --- | --- | --- | --- | --- | --- | --- | --- | --- | --- | --- |
|  |  | **Average model** |  | **Basal area** | |  | **Precipitation** | |  | **Temperature** | |
|  |  |  |  | Low | High |  | Low | High |  | Low | High |
| 10 |  | 10.9 |  | 10.6 | 11.1 |  | 10.5 | 11.2 |  | 11.5 | 10.3 |
| 15 |  | 12.7 |  | 12.3 | 13 |  | 12.2 | 13.1 |  | 13.2 | 12.2 |
| 20 |  | 14.2 |  | 13.7 | 14.6 |  | 13.6 | 14.7 |  | 14.6 | 13.7 |
| 25 |  | 15.4 |  | 14.9 | 15.9 |  | 14.8 | 16.0 |  | 15.7 | 15.1 |
| 30 |  | **16.5** |  | **15.9** | **17.1** |  | **15.8** | **17.2** |  | **16.8** | **16.3** |
| 40 |  | 18.4 |  | 17.7 | 19.2 |  | 17.6 | 19.3 |  | 18.5 | 18.3 |
| 50 |  | 20.1 |  | 19.2 | 21.0 |  | 19.1 | 21.1 |  | 20.0 | 20.1 |
| 60 |  | 21.5 |  | 20.6 | 22.5 |  | 20.5 | 22.6 |  | 21.3 | 21.7 |
| 70 |  | 22.8 |  | 21.8 | 23.9 |  | 21.7 | 24.1 |  | 22.5 | 23.2 |
| 80 |  | 24.0 |  | 22.9 | 25.2 |  | 22.7 | 25.4 |  | 23.6 | 24.5 |
| 90 |  | 25.1 |  | 23.9 | 26.4 |  | 23.8 | 26.6 |  | 24.5 | 25.7 |
| 100 |  | 26.2 |  | 24.8 | 27.5 |  | 24.7 | 27.7 |  | 25.4 | 26.9 |

**Table S6:** Predicted tree crown radius, *Crad*, at different values of *D* using the fitted model and considering low and high basal area; low and high precipitation; and low and high temperature. Predictions of *Crad* in the table are based on equation 3 in the main text. Average model predictions are based on 0 effect sizes for all the predictors. Low and high represent one-standard deviation decrease (-1*mean effect size) and increase (+1*mean effect size) respectively, for basal area, precipitation and temperature.

| *D* (cm) |  | Predicted *Crad* (m) | | | | | | | | | |
| --- | --- | --- | --- | --- | --- | --- | --- | --- | --- | --- | --- |
|  |  | **Average model** |  | **Basal area** | |  | **Precipitation** | |  | **Temperature** | |
|  |  |  |  | Low | High |  | Low | High |  | Low | High |
| 10 |  | 1.9 |  | 2.1 | 1.8 |  | 2.0 | 1.9 |  | 2.0 | 1.9 |
| 15 |  | 2.7 |  | 2.8 | 2.6 |  | 2.7 | 2.6 |  | 2.7 | 2.7 |
| 20 |  | 3.4 |  | 3.5 | 3.2 |  | 3.4 | 3.3 |  | 3.3 | 3.4 |
| 25 |  | 4.0 |  | 4.1 | 3.9 |  | 4.0 | 4.0 |  | 3.9 | 4.1 |
| 30 |  | **4.6** |  | **4.7** | **4.5** |  | **4.6** | **4.6** |  | **4.5** | **4.8** |
| 40 |  | 5.8 |  | 5.8 | 5.8 |  | 5.8 | 5.8 |  | 5.5 | 6.1 |
| 50 |  | 6.9 |  | 6.9 | 6.9 |  | 6.9 | 6.9 |  | 6.5 | 7.3 |
| 60 |  | 8.0 |  | 7.9 | 8.0 |  | 7.9 | 8.0 |  | 7.4 | 8.6 |
| 70 |  | 9.0 |  | 8.8 | 9.1 |  | 9.0 | 9.0 |  | 8.3 | 9.7 |
| 80 |  | 10.0 |  | 9.8 | 10.2 |  | 9.9 | 10.0 |  | 9.2 | 10.9 |
| 90 |  | 10.9 |  | 10.7 | 11.2 |  | 10.9 | 11.0 |  | 10.0 | 12.0 |
| 100 |  | 11.9 |  | 11.5 | 12.3 |  | 11.8 | 12.0 |  | 10.8 | 13.1 |

**Table S7:** Predicted tree crown depth, *Cdep*, at different values of *D* using the fitted model and considering low and high basal area; low and high precipitation; and low and high temperature. Predictions of *Cdep* in the table are based on equation 3 in the main text. Average model predictions are based on 0 effect sizes for all the predictors. Low and high represent one-standard deviation decrease (-1*mean effect size) and increase (+1*mean effect size) respectively, for basal area, precipitation and temperature.

| *D* (cm) |  | Predicted *Cdep* (m) | | | | | | | | | |
| --- | --- | --- | --- | --- | --- | --- | --- | --- | --- | --- | --- |
|  |  | **Average model** |  | **Basal area** | |  | **Precipitation** | |  | **Temperature** | |
|  |  |  |  | Low | High |  | Low | High |  | Low | High |
| 10 |  | 4.0 |  | 4.1 | 3.9 |  | 4.0 | 4.0 |  | 4.0 | 4.1 |
| 15 |  | 4.6 |  | 4.6 | 4.5 |  | 4.4 | 4.7 |  | 4.2 | 4.9 |
| 20 |  | 5.0 |  | 5.0 | 5.0 |  | 4.7 | 5.3 |  | 4.4 | 5.6 |
| 25 |  | 5.3 |  | 5.3 | 5.3 |  | 4.9 | 5.8 |  | 4.5 | 6.3 |
| 30 |  | **5.6** |  | **5.5** | **5.7** |  | **5.1** | **6.2** |  | **4.6** | **6.8** |
| 40 |  | 6.1 |  | 6.0 | 6.2 |  | 5.4 | 6.9 |  | 4.8 | 7.8 |
| 50 |  | 6.5 |  | 6.3 | 6.7 |  | 5.7 | 7.5 |  | 4.9 | 8.7 |
| 60 |  | 6.9 |  | 6.7 | 7.1 |  | 5.9 | 8.1 |  | 5.0 | 9.5 |
| 70 |  | 7.2 |  | 6.9 | 7.5 |  | 6.1 | 8.6 |  | 5.1 | 10.2 |
| 80 |  | 7.5 |  | 7.2 | 7.9 |  | 6.3 | 9.0 |  | 5.2 | 10.9 |
| 90 |  | 7.8 |  | 7.4 | 8.2 |  | 6.4 | 9.5 |  | 5.3 | 11.5 |
| 100 |  | 8.0 |  | 7.6 | 8.5 |  | 6.6 | 9.9 |  | 5.3 | 12.1 |

**Table S8:** Estimated mean [95% credible interval] of the model parameters, $\alpha$ and $\beta$ , and effect sizes of basal area, precipitation and temperature on $\alpha$ and $\beta$ of the power law model for the NFI dataset for inter- and intra-specific *H*-*D* allometry. Significant (excluding infinitesimal and marginal – where |0.00| is the lower and/or upper bound of the 95% credible interval) parameters and effect sizes are shown in bold. $\alpha$ estimates for all relationships and variables are on natural log scale. The effect size for each predictor variable represents the change in $\alpha$ and/or $\beta$ as a result of one-standard deviation increase in the predictor variable.

| Relationship | **Variable** |  | **Model parameter** | | |
| --- | --- | --- | --- | --- | --- |
|  |  |  | $\alpha$ Estimate [95% CI] |  | $\beta$ Estimate [95% CI] |
| *H-D* | **Intercept** |  | **0.93 [0.90, 0.96]** |  | **0.39 [0.38, 0.40]** |
| *Inter-specific effect size* | | | | | |
|  | **Basal area** |  | **0.10 [0.05, 0.14]** |  | 0.00 [0.00, 0.02] |
|  | Precipitation |  | -0.05 [-0.12, -0.00] |  | 0.01 [0.00, 0.03] |
|  | Temperature |  | 0.02 [-0.04, 0.07] |  | 0.01 [0.00, 0.03] |
| *Intra-specific effect size* | | | | | |
|  | **Basal area** |  | **0.10 [0.08, 0.11]** |  | 0.00 [0.00, 0.01] |
|  | **Precipitation** |  | **-0.04 [-0.06, -0.02]** |  | 0.01 [0.00, 0.02] |
|  | **Temperature** |  | **0.03 [0.01, 0.04]** |  | 0.00 [0.00, 0.01] |

**Table S9:** Predicted tree height, *H*, at different values of *D* using the fitted model, equation 4 in the main text, and considering low and high basal area. Average model predictions are based on 0 effect size for basal area. Low and high represent one-standard deviation decrease (-1*mean effect size) and increase (+1*mean effect size) respectively, for basal area.

| *D* (cm) | **Predicted *H* (m) for tall-stature species** | | | | | | | | | | | |
| --- | --- | --- | --- | --- | --- | --- | --- | --- | --- | --- | --- | --- |
|  | *Brachystegia boehmii* | | | *Brachystegia longifolia* | | | *Brachystegia spiciformis* | | | *Julbernardia paniculata* | | |
|  | Average model | Low | High | Average model | Low | High | Average model | Low | High | Average model | Low | High |
| 10 | 7.1 | 6.3 | 8.1 | 7.2 | 6.4 | 8.1 | 7.9 | 7.0 | 8.9 | 7.8 | 6.9 | 8.8 |
| 15 | 8.4 | 7.4 | 9.5 | 8.6 | 7.6 | 9.7 | 9.2 | 8.1 | 10.4 | 9.1 | 8.0 | 10.3 |
| 20 | 9.4 | 8.3 | 10.6 | 9.7 | 8.6 | 10.9 | 10.2 | 9.1 | 11.6 | 10.2 | 9.0 | 11.5 |
| 25 | 10.2 | 9.0 | 11.6 | 10.6 | 9.4 | 12.0 | 11.1 | 9.8 | 12.6 | 11.1 | 9.8 | 12.6 |
| 30 | **11.0** | **9.7** | **12.5** | **11.5** | **10.2** | **13.0** | **11.9** | **10.6** | **13.5** | **11.9** | **10.5** | **13.5** |
| 40 | 12.3 | 10.9 | 14.0 | 13.0 | 11.5 | 14.7 | 13.3 | 11.8 | 15.1 | 13.3 | 11.7 | 15.0 |
| 50 | 13.5 | 11.9 | 15.3 | 14.3 | 12.6 | 16.2 | 14.5 | 12.8 | 16.4 | 14.4 | 12.7 | 16.4 |
| 60 | 14.5 | 12.7 | 16.5 | 15.4 | 13.6 | 17.5 | 15.5 | 13.7 | 17.6 | 15.5 | 13.6 | 17.6 |
| 70 | 15.4 | 13.5 | 17.6 | 16.5 | 14.5 | 18.7 | 16.5 | 14.5 | 18.6 | 16.4 | 14.5 | 18.6 |
| 80 | 16.3 | 14.3 | 18.5 | 17.4 | 15.3 | 19.8 | 17.3 | 15.3 | 19.6 | 17.3 | 15.2 | 19.6 |
| 90 | 17.0 | 14.9 | 19.4 | 18.3 | 16.1 | 20.8 | 18.1 | 16.0 | 20.5 | 18.1 | 15.9 | 20.5 |
| 100 | 17.8 | 15.6 | 20.3 | 19.2 | 16.9 | 21.8 | 18.8 | 16.6 | 21.4 | 18.8 | 16.6 | 21.4 |

**Table S10:** Predicted tree height, *H*, at different values of *D* using the fitted model, equation 4 in the main text, and considering low and high basal area. Average model predictions are based on 0 effect size for basal area. Low and high represent one-standard deviation decrease (-1*mean effect size) and increase (+1*mean effect size) respectively, for basal area.

| *D* (cm) | **Predicted *H* (m) for short-stature species** | | | | | | | | | | | |
| --- | --- | --- | --- | --- | --- | --- | --- | --- | --- | --- | --- | --- |
|  | *Diplorhynchus condylocarpon* | | | *Pseudolachnostylis maprouneifolia* | | | *Uapaca kirkiana* | | | *Monotes africanus* | | |
|  | Average model | Low | High | Average model | Low | High | Average model | Low | High | Average model | Low | High |
| 10 | 6.6 | 6.1 | 7.2 | 6.1 | 5.6 | 6.6 | 6.4 | 5.7 | 7.3 | 6.6 | 6.0 | 7.4 |
| 15 | 7.7 | 7.1 | 8.3 | 7.2 | 6.6 | 7.8 | 7.5 | 6.6 | 8.5 | 7.9 | 7.1 | 8.7 |
| 20 | 8.5 | 7.9 | 9.2 | 8.0 | 7.4 | 8.7 | 8.4 | 7.4 | 9.5 | 8.9 | 8.0 | 9.9 |
| 25 | 9.3 | 8.5 | 10.0 | 8.8 | 8.1 | 9.5 | 9.1 | 8.0 | 10.3 | 9.7 | 8.7 | 10.8 |
| 30 | **9.9** | **9.1** | **10.7** | **9.4** | **8.7** | **10.2** | **9.7** | **8.6** | **11.0** | **10.5** | **9.4** | **11.7** |
| 40 | 11.0 | 10.1 | 11.9 | 10.6 | 9.7 | 11.5 | 10.9 | 9.6 | 12.3 | 11.8 | 10.6 | 13.2 |
| 50 | 11.9 | 11.0 | 12.9 | 11.5 | 10.6 | 12.5 | 11.8 | 10.4 | 13.4 | 13.0 | 11.6 | 14.5 |
| 60 | 12.7 | 11.8 | 13.8 | 12.4 | 11.4 | 13.5 | 12.7 | 11.1 | 14.4 | 14.0 | 12.5 | 15.6 |
| 70 | 13.5 | 12.4 | 14.6 | 13.1 | 12.1 | 14.3 | 13.4 | 11.8 | 15.2 | 14.9 | 13.4 | 16.7 |
| 80 | 14.1 | 13.1 | 15.3 | 13.9 | 12.7 | 15.1 | 14.1 | 12.4 | 16.0 | 15.8 | 14.1 | 17.6 |
| 90 | 14.8 | 13.6 | 16.0 | 14.5 | 13.3 | 15.8 | 14.8 | 13.0 | 16.8 | 16.6 | 14.8 | 18.5 |
| 100 | 15.3 | 14.2 | 16.6 | 15.1 | 13.9 | 16.5 | 15.3 | 13.5 | 17.5 | 17.3 | 15.5 | 19.4 |

### Supplementary Figures

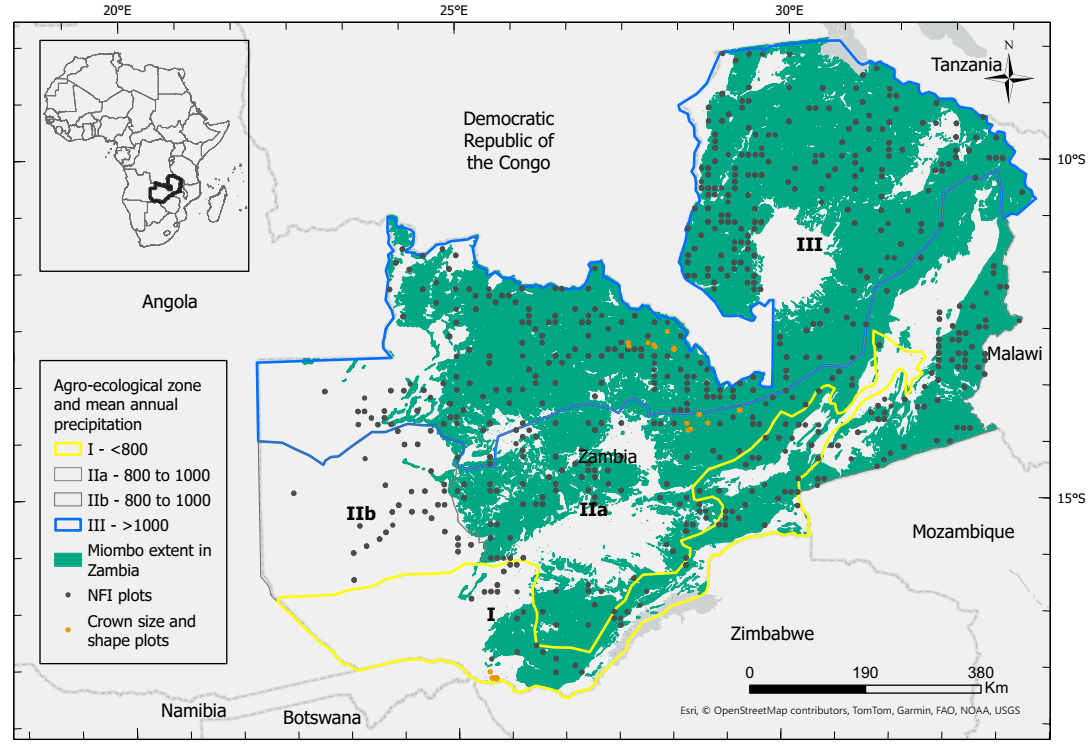

**Fig. S1:** The location of NFI and crown size and shape plots in Zambia.

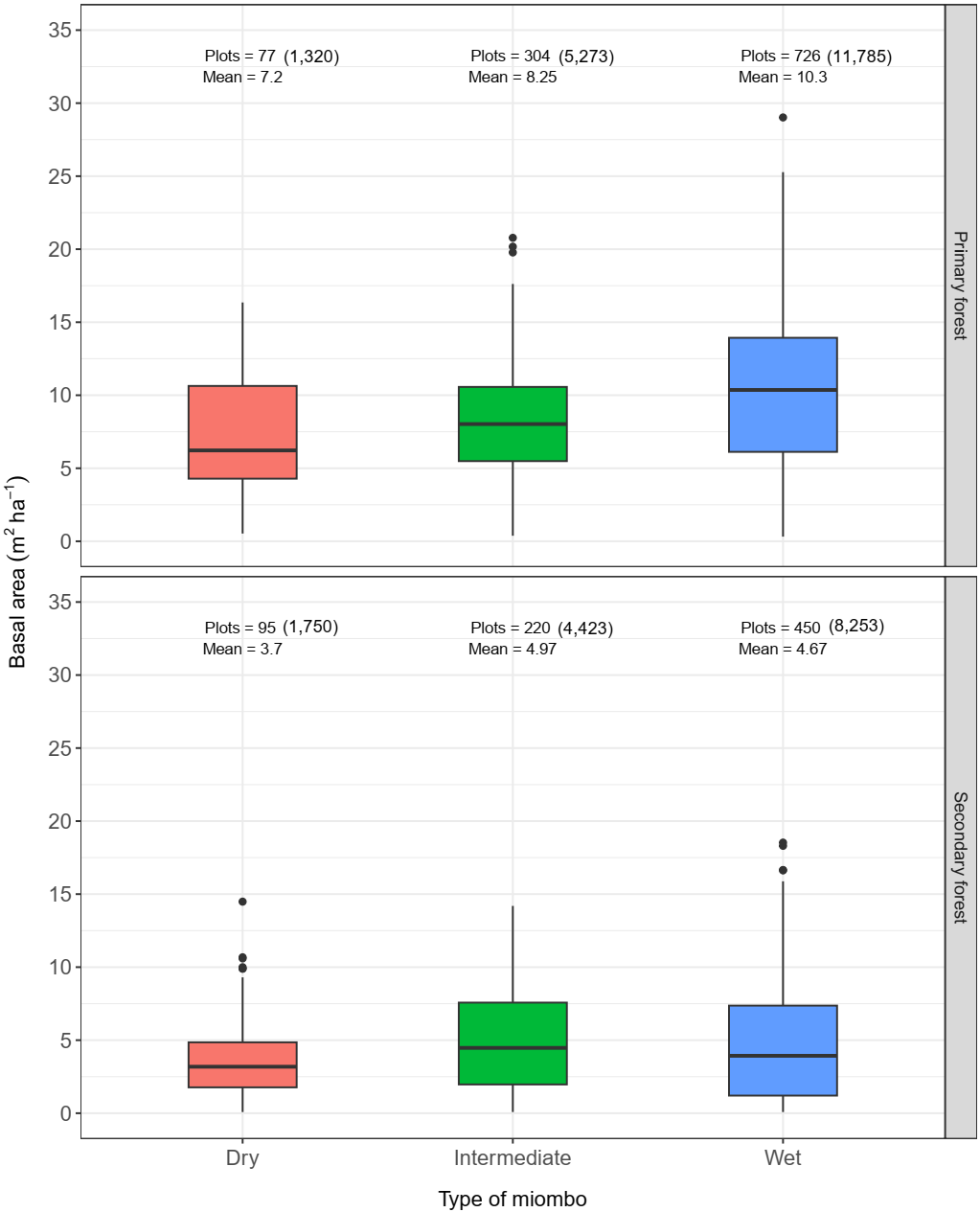

**Fig. S2:** Basal area in NFI plots classified as primary and secondary forest in three types of miombo. The number of trees in plots under different categories are shown in brackets beside number of plots

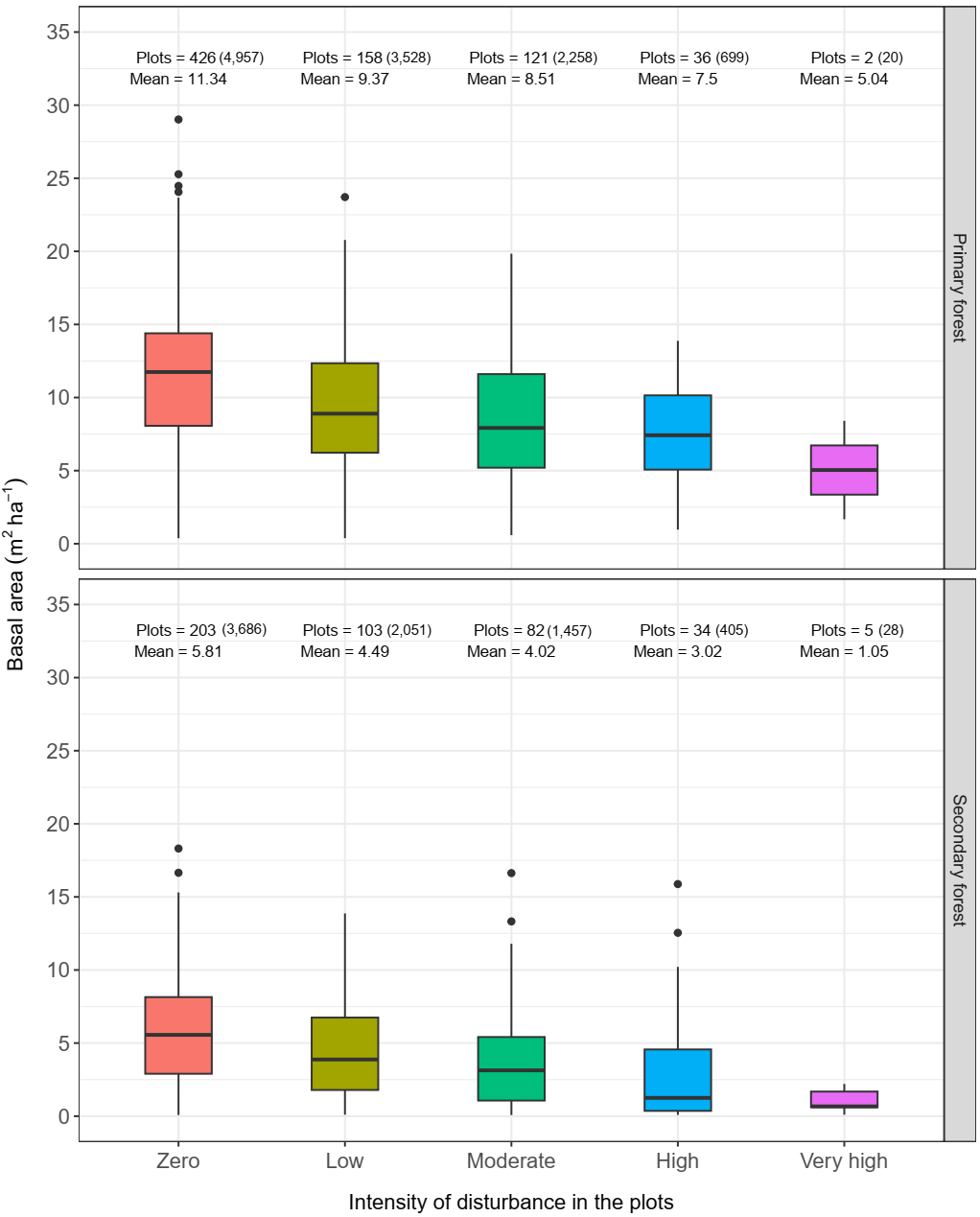

**Fig. S3:** Basal area versus intensity of disturbance in plots classified as primary forest and secondary forest for the NFI dataset. The number of trees in plots under different categories are shown in brackets beside number of plots

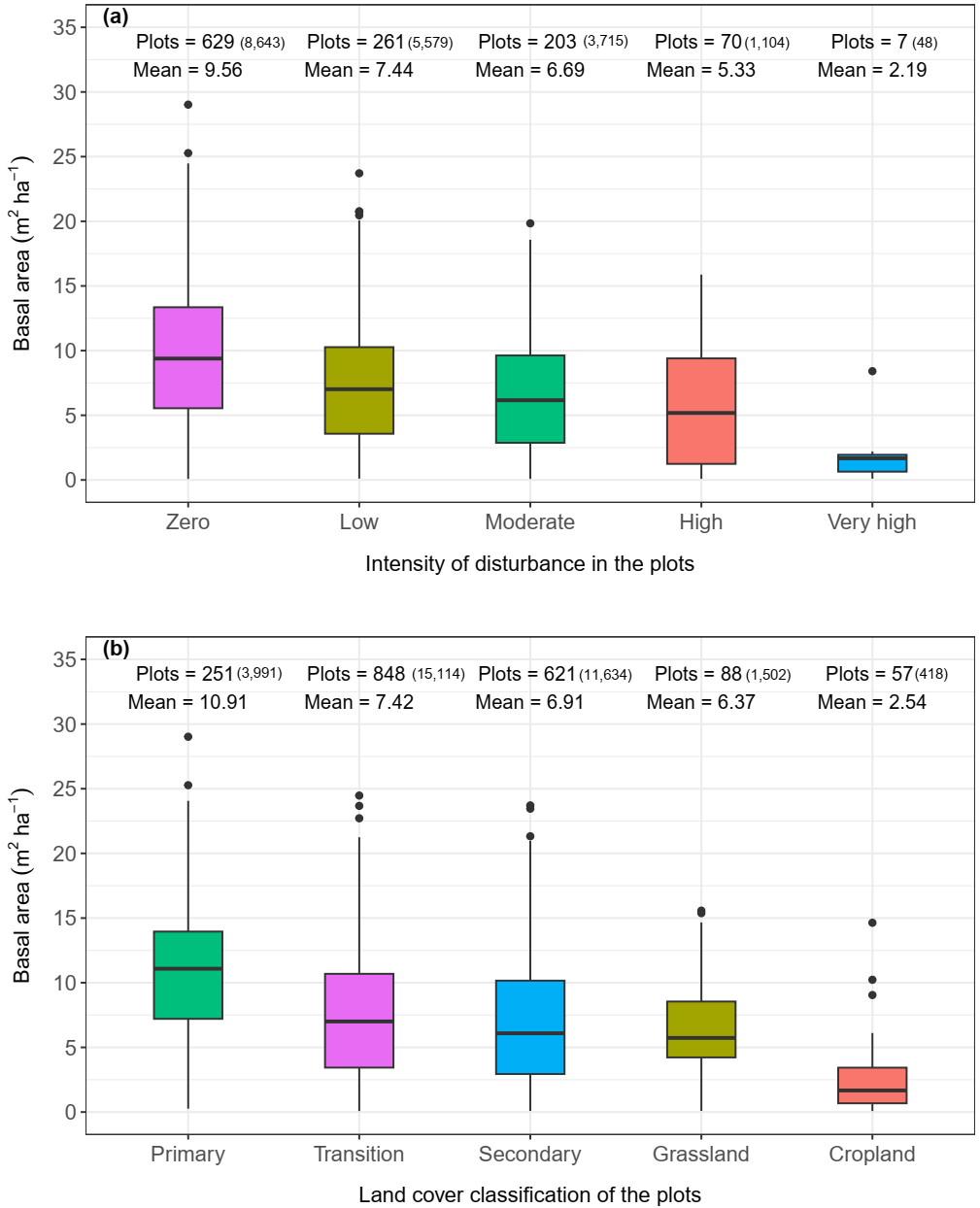

**Fig. S4:** Basal area versus (a) intensity of disturbance for plots with zero, one or two or three of the problems (burning, overexploitation of resources and overgrazing) identified per plot in the NFI dataset; and (b) land cover classification based on the works of (Phiri et al., 2019). The number of trees in plots under different categories are shown in brackets beside number of plots.

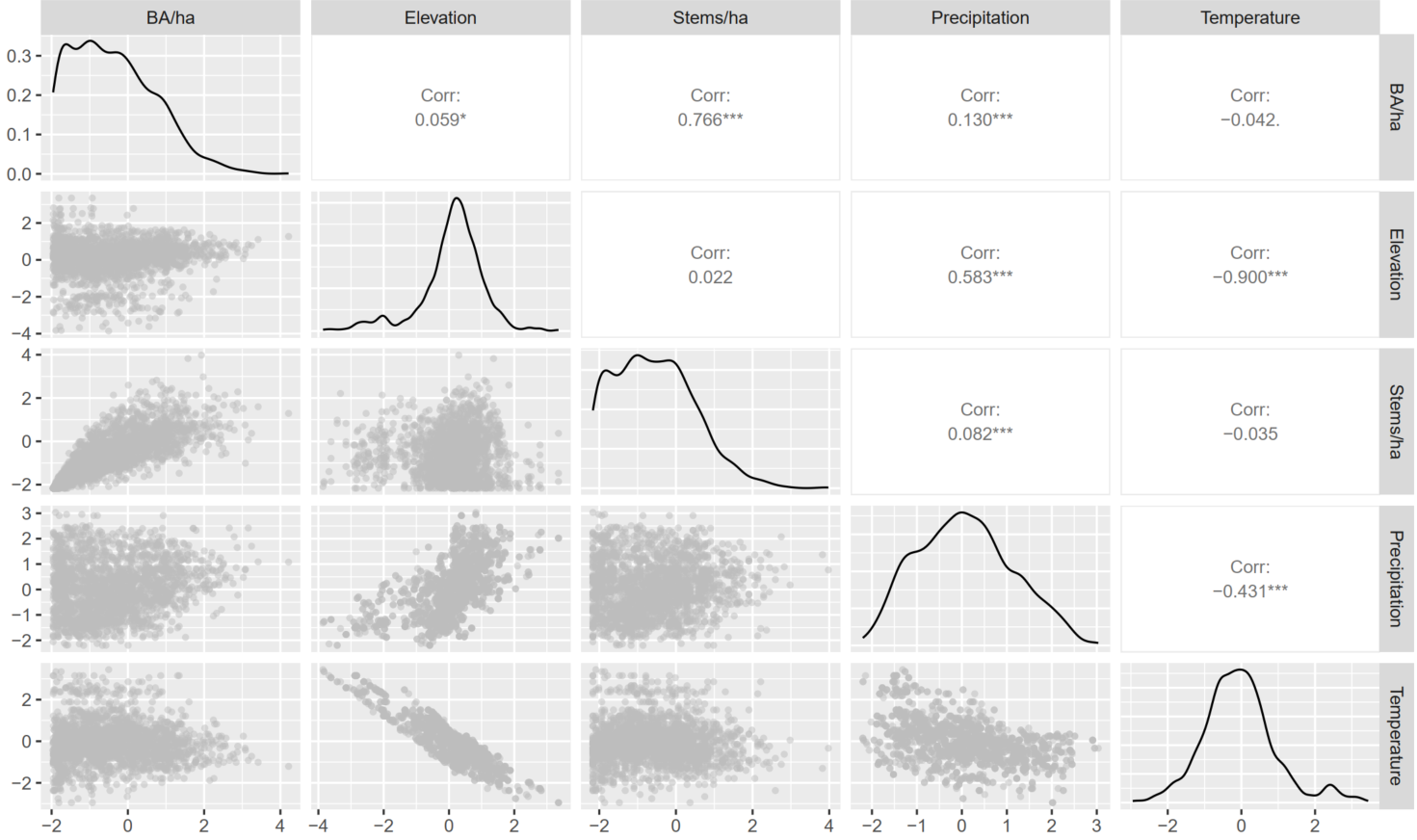

**Fig. S5:** Correlation matrix among key variables (scaled) considered in this study. BA/ha is basal area per hectare. The variables were calculated at plot level.

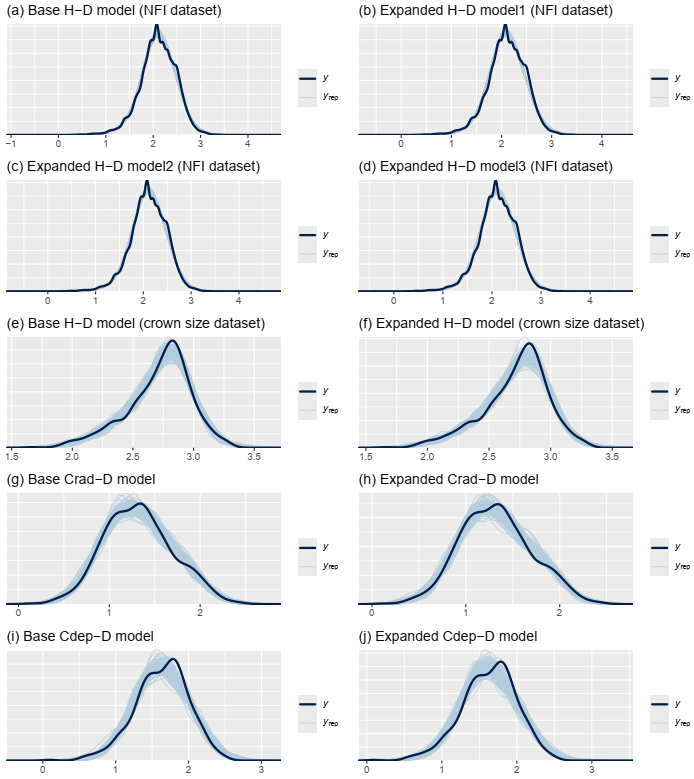

**Fig. S6:** Posterior predictive checks for the fitted *H*-*D* and crown size models. The dark blue curve ($y$) represents log transformed values of observed tree *H*, *Crad* and *Cdep* while the light blue curve/lines represents 100 new samples ($y_{rep}$) of predicted tree *H*, *Crad* and *Cdep* based on posterior distribution of parameters for each model. Model fits are considered adequate if the two distributions, i.e. $y$ and $y_{rep}$, look similar or overlap. Expanded H-D model1 is based on equation 3 while expanded H-D model2 and model3 are based on equation 4 in the main text.

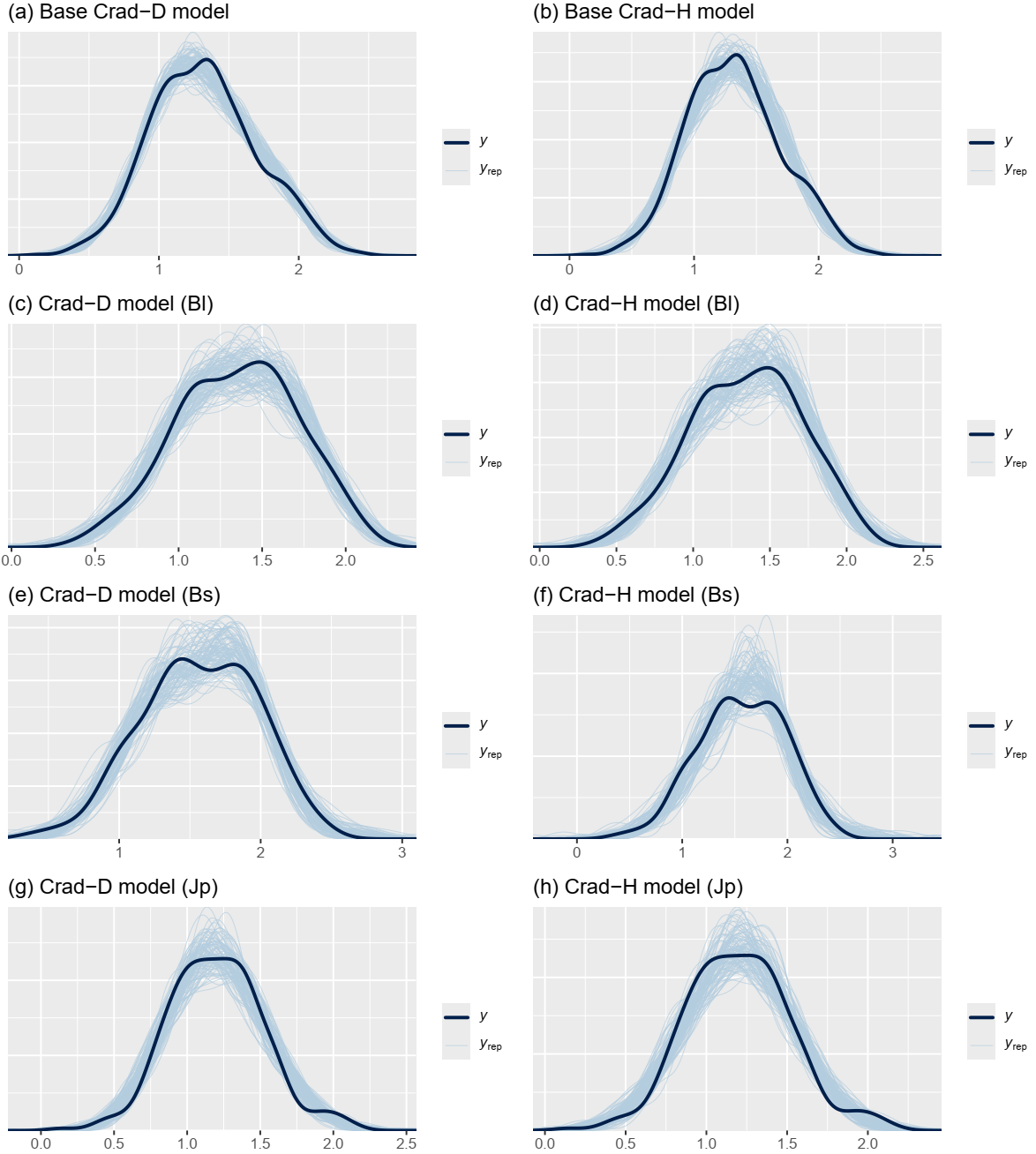

**Fig. S7:** Posterior predictive checks for the fitted base crown size models across species and for each of three species - *Brachystegia longifolia* (Bl) , *Brachystegia spiciformis* (Bs) and *Julbernardia paniculata* (Jp). The dark blue curve ($y$) represents log transformed values of observed tree *Crad* and *Cdep* while the light blue curve/lines represents 100 new samples ($y_{rep}$) of predicted tree *Crad* and *Cdep* based on posterior distribution of parameters for each model. Model fits are considered adequate if the two distributions, i.e. $y$ and $y_{rep}$, look similar or overlap.

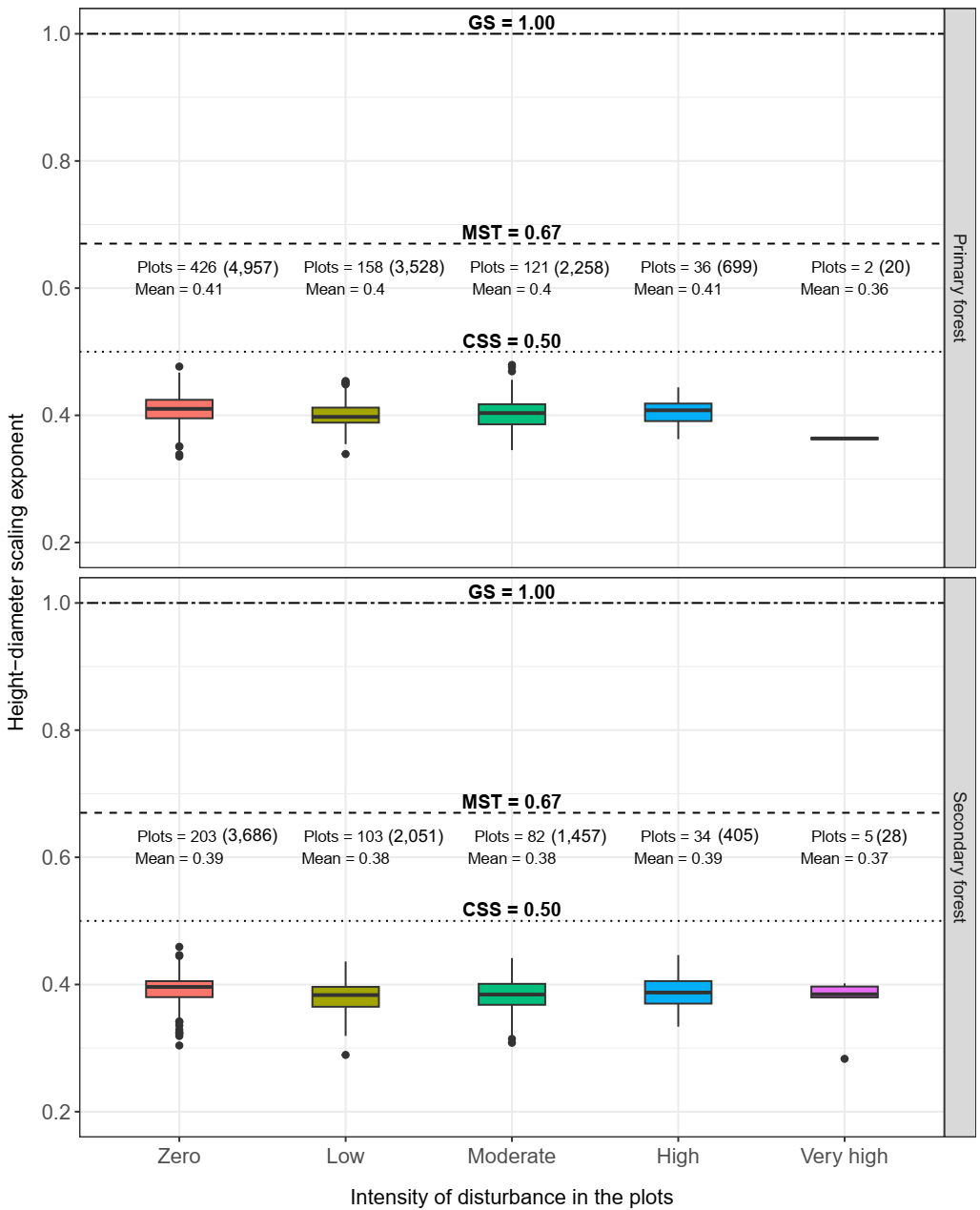

**Fig. S8:** Variation in estimated scaling exponents for *H-D* allometric relationship in primary and secondary forests and across different intensity of disturbance (zero, low, high and very high). Theoretical predictions of MST (dashed; 0.67), CSS (dotted; 0.50) and GS (two dash; 1.00) are shown as horizontal lines in each panel. The number of trees in plots under different categories are shown in brackets beside number of plots.

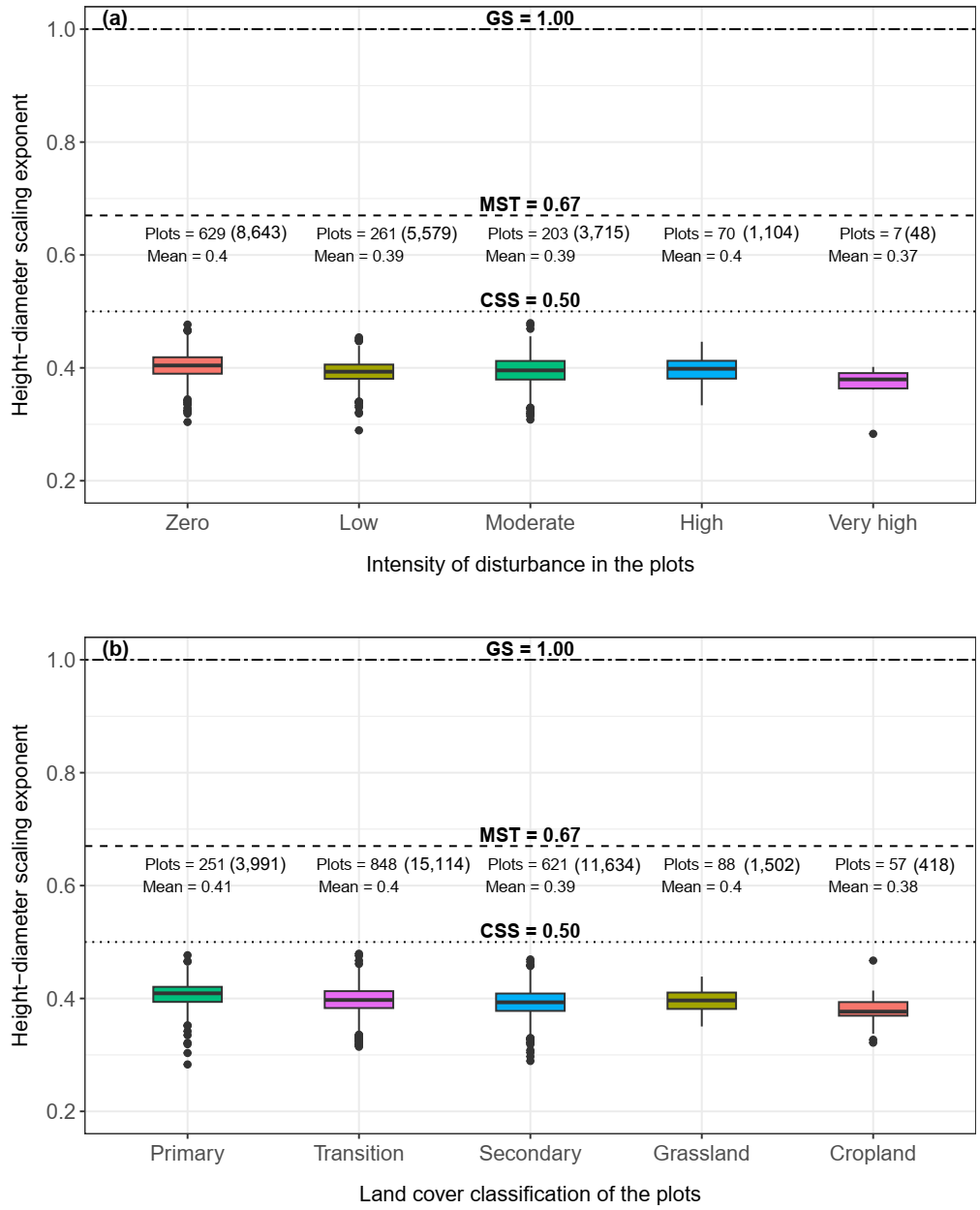

**Fig. S9:** Variation in estimated scaling exponents for *H-D* allometric relationship in **(a)** different intensity of disturbance (zero, low, high and very high) for plots with zero or one or two or three of the problems (burning, overexploitation of resources and overgrazing) identified per plot in the NFI dataset; and **(b)** land cover classification. Theoretical predictions of MST (dashed; 0.67), CSS (dotted; 0.50) and GS (two dash; 1.00) are shown as horizontal lines in each panel. The number of trees in plots under different categories are shown in brackets beside number of plots.

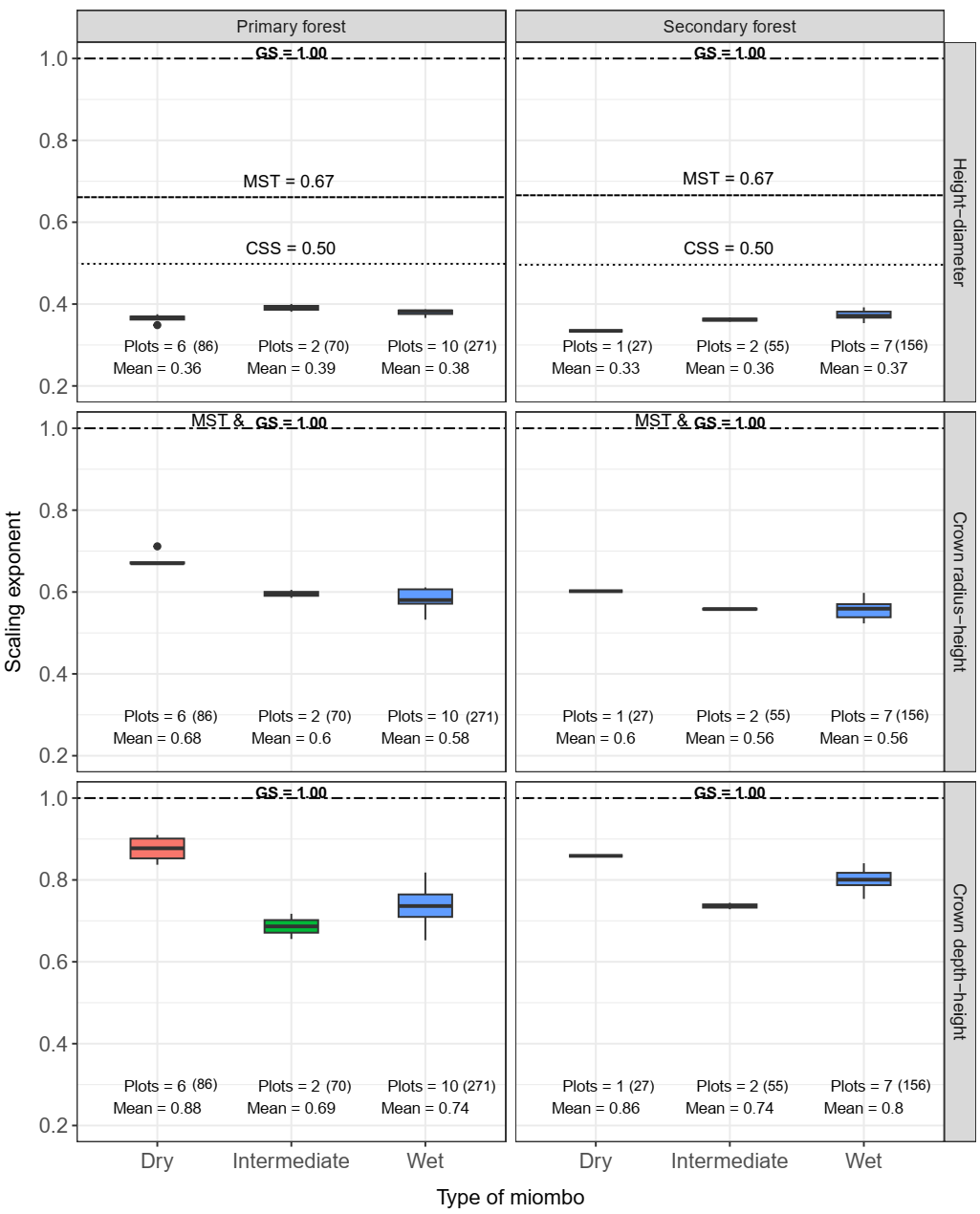

**Fig. S10:** Variation in estimated scaling exponents for different allometric relationships in primary and secondary forests and across three types of miombo (dry, intermediate and wet) for the crown size and shape dataset. Theoretical predictions of MST (dashed; 0.67), CSS (dotted; 0.50) and GS (two dash; 1.00) are shown as horizontal lines in each panel. The number of trees in plots under different categories are shown in brackets beside number of plots.

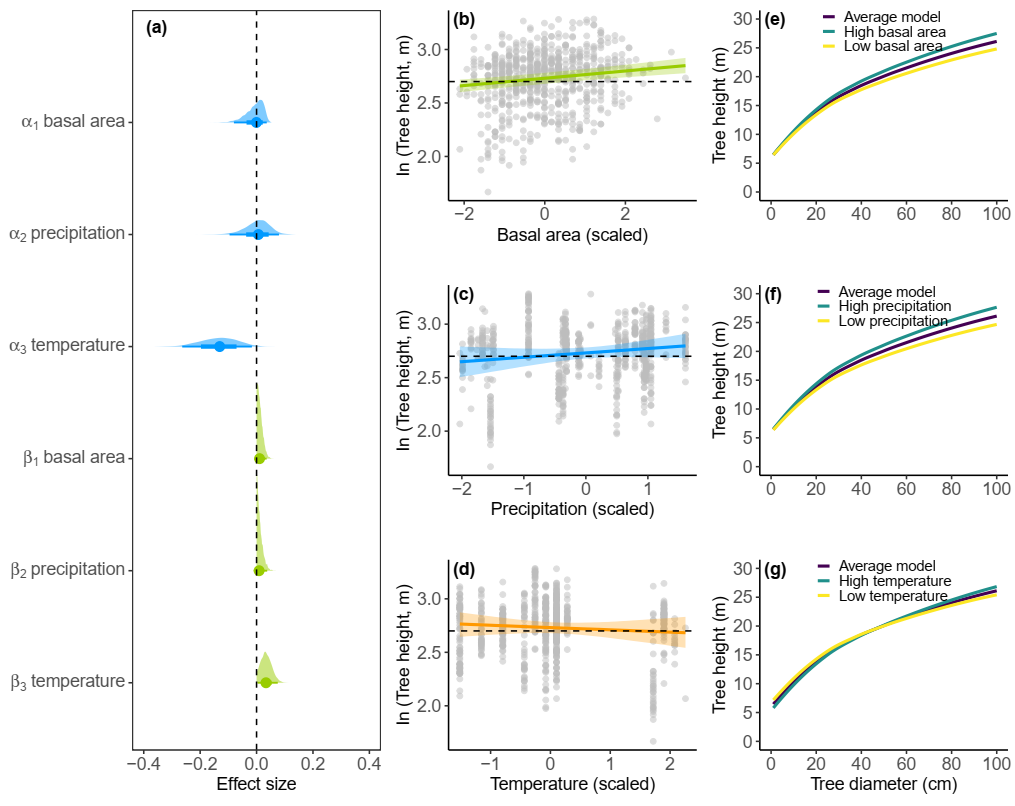

**Fig. S11:** Effects of basal area, precipitation and temperature on **(a)** the height–diameter (*H-D*) power law model parameters, **(b**–**d)** *H* predictions for a tree of mean size along gradients of basal area, precipitation and temperature, and (**e**–**g**) *H* predictions across a range of *D* values (1-100 cm) while varying one model predictor (e.g., basal area in **e**) and keeping the other two (precipitation and temperature) constant at their mean value and setting their effects to zero. In (**b**–**d**), coloured lines represent mean model predictions (with shaded 95% credible intervals), the horizontal dashed line represents the mean tree *H* (on log scale) across the crown size and shape dataset, and data points are shown as grey circles. In (**e**–**g**) low (yellow line) and high (blue line) prediction scenarios correspond to ± 1 standard deviation.

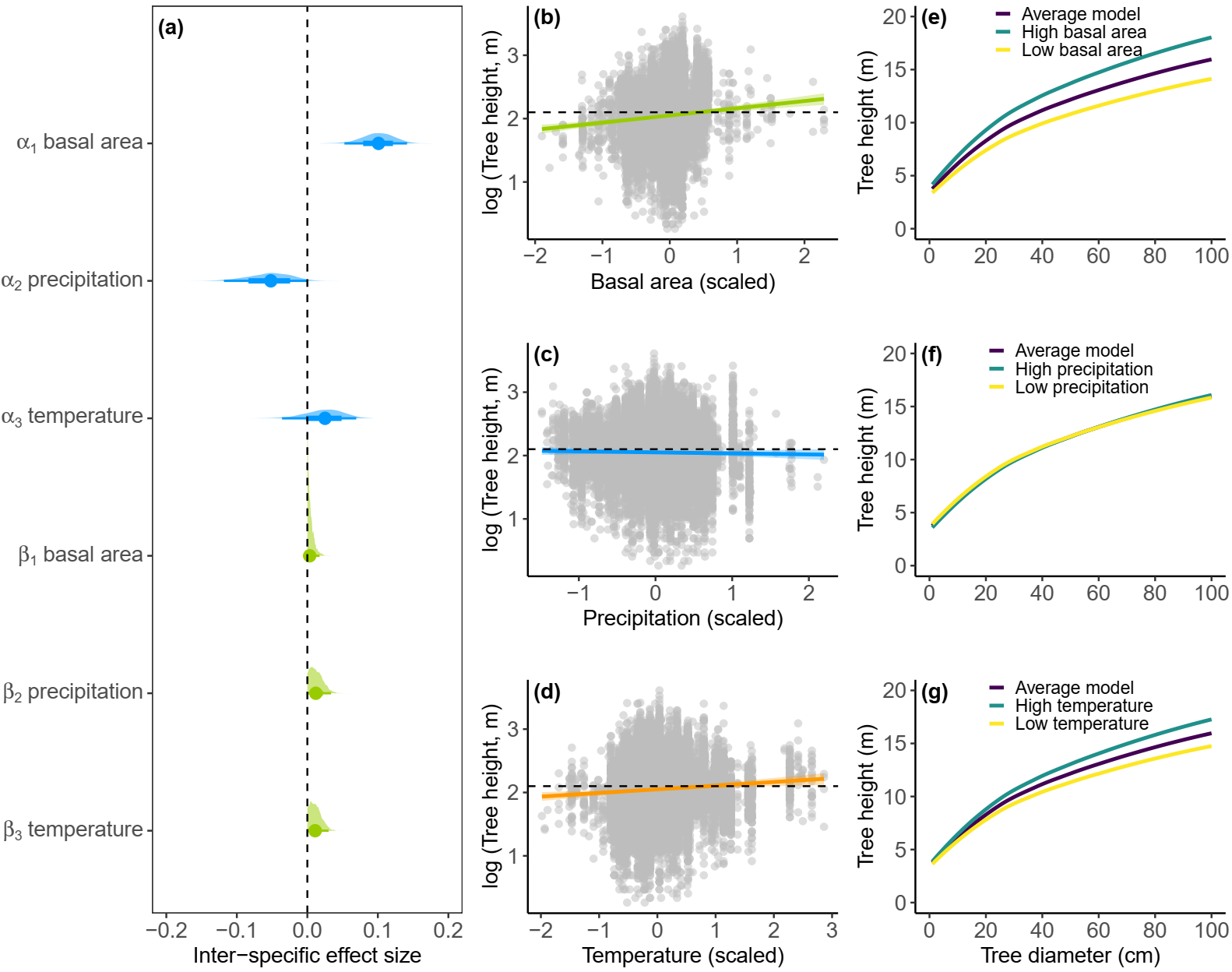

**Fig. S12:** Effects (inter-specific) of basal area, precipitation and temperature on **(a)** the height–diameter (*H-D*) power law model parameters, **(b**–**d)** *H* predictions for a tree of mean size along gradients of basal area, precipitation and temperature, and (**e**–**g**) *H* predictions across a range of *D* values (1-100 cm) while varying one model predictor (e.g., basal area in **e**) and keeping the other two (precipitation and temperature) constant at their mean value and setting their effects to zero. In (**b**–**d**), coloured lines represent mean model predictions (with shaded 95% credible intervals), the horizontal dashed line represents the mean tree *H* (on log scale) across the NFI dataset, and data points are shown as grey circles. In (**e**–**g**) low (yellow line) and high (blue line) prediction scenarios correspond to ± 1 standard deviation.

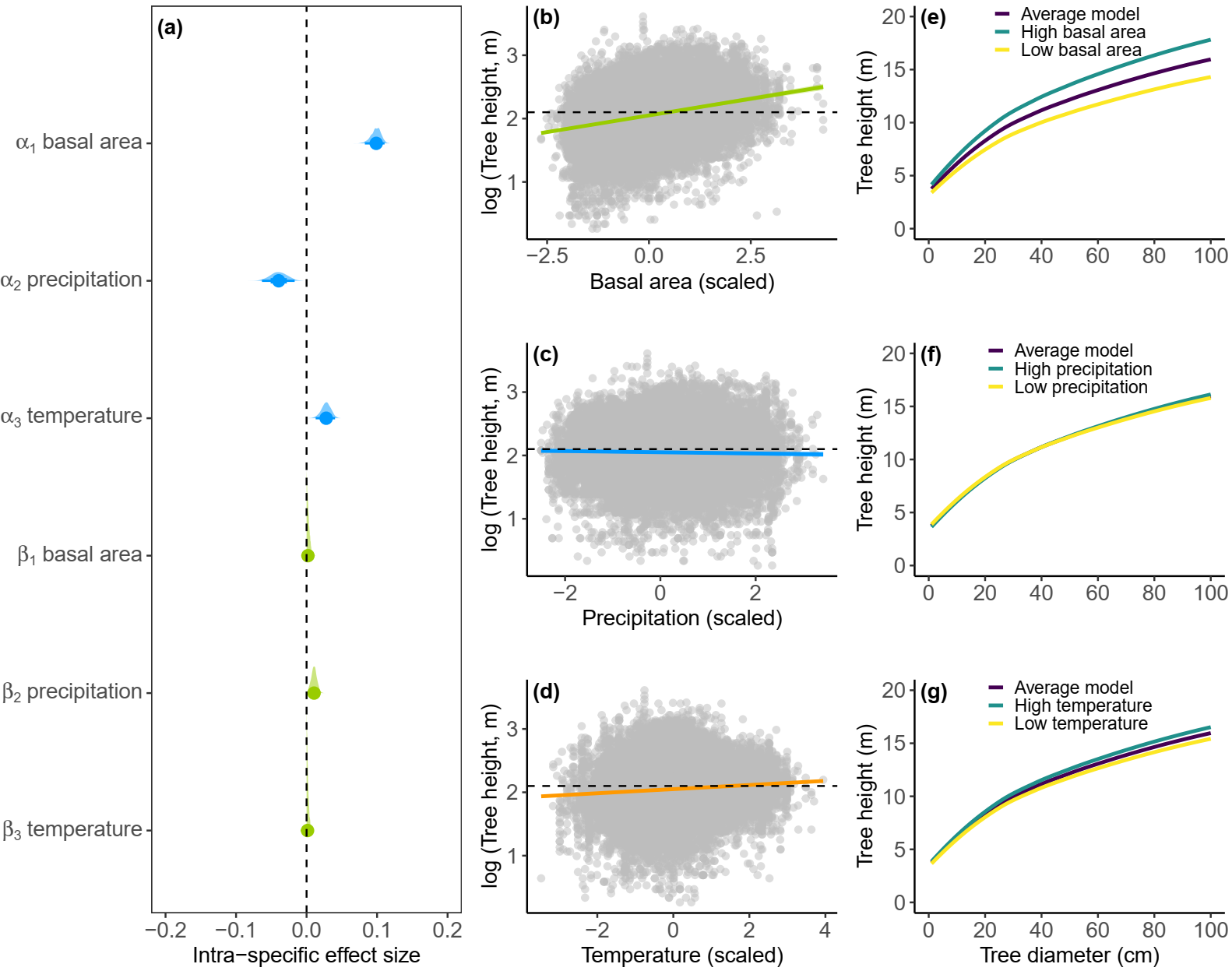

**Fig. S13:** Effects (intra-specific) of basal area, precipitation and temperature on **(a)** the height–diameter (*H-D*) power law model parameters, **(b**–**d)** *H* predictions for a tree of mean size along gradients of basal area, precipitation and temperature, and (**e**–**g**) *H* predictions across a range of *D* values (1-100 cm) while varying one model predictor (e.g., basal area in **e**) and keeping the other two (precipitation and temperature) constant at their mean value and setting their effects to zero. In (**b**–**d**), coloured lines represent mean model predictions (with shaded 95% credible intervals), the horizontal dashed line represents the mean tree *H* (on log scale) across the NFI dataset, and data points are shown as grey circles. In (**e**–**g**) low (yellow line) and high (blue line) prediction scenarios correspond to ± 1 standard deviation.

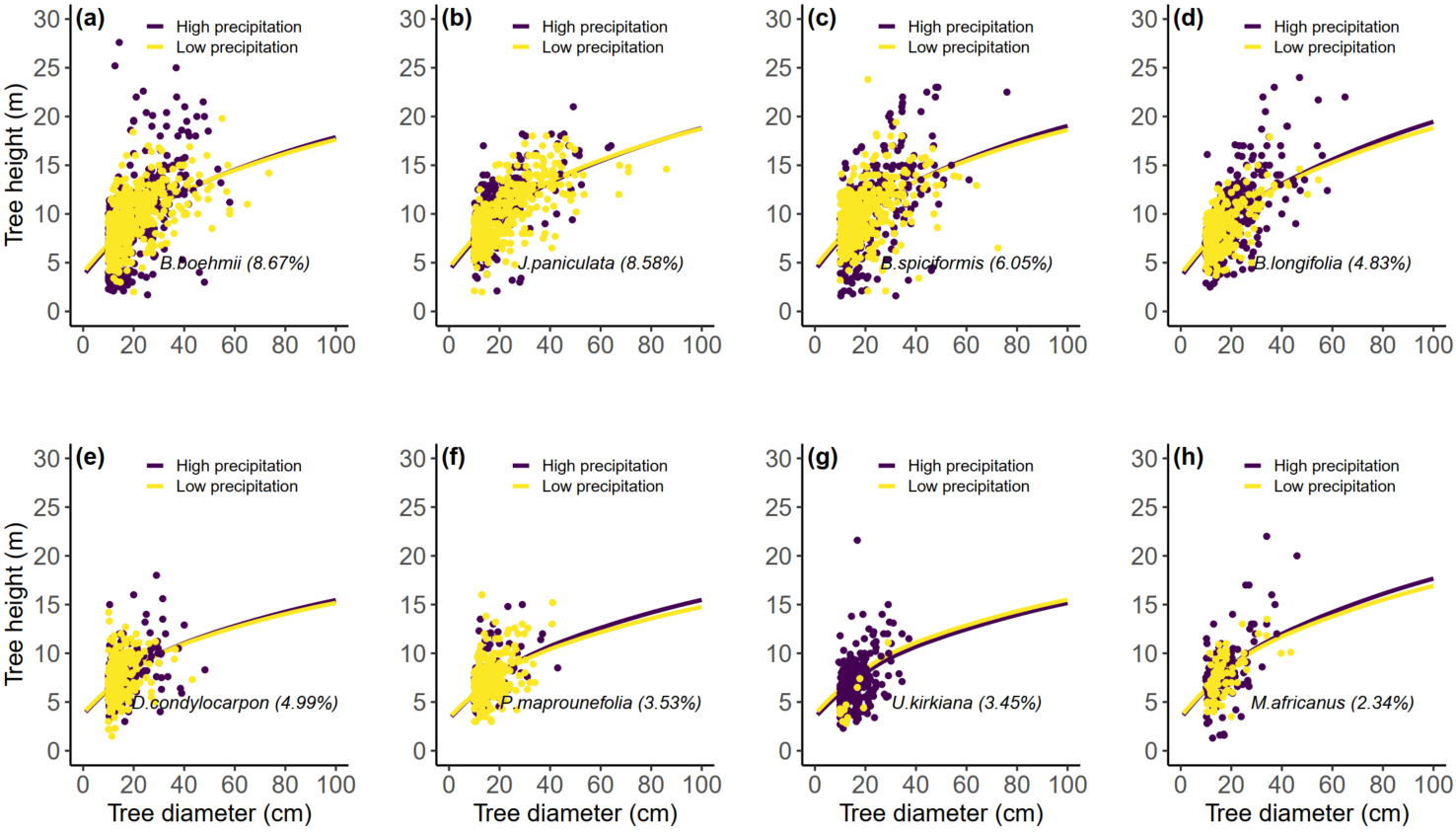

**Fig. S14:** Effects of precipitation on intra-specific variability in *H-D* scaling relationships for the four most abundant tall (**a**–**d**) and short (**e**–**h**) stature miombo tree species. Lines represent predictions for one-standard deviation increase (purple) and decrease (yellow) in precipitation. Circles represent raw data from plots that are either one-standard deviation above (purple) or below (yellow) mean precipitation. The percentage in brackets beside each species name corresponds to its relative abundance in the NFI data.

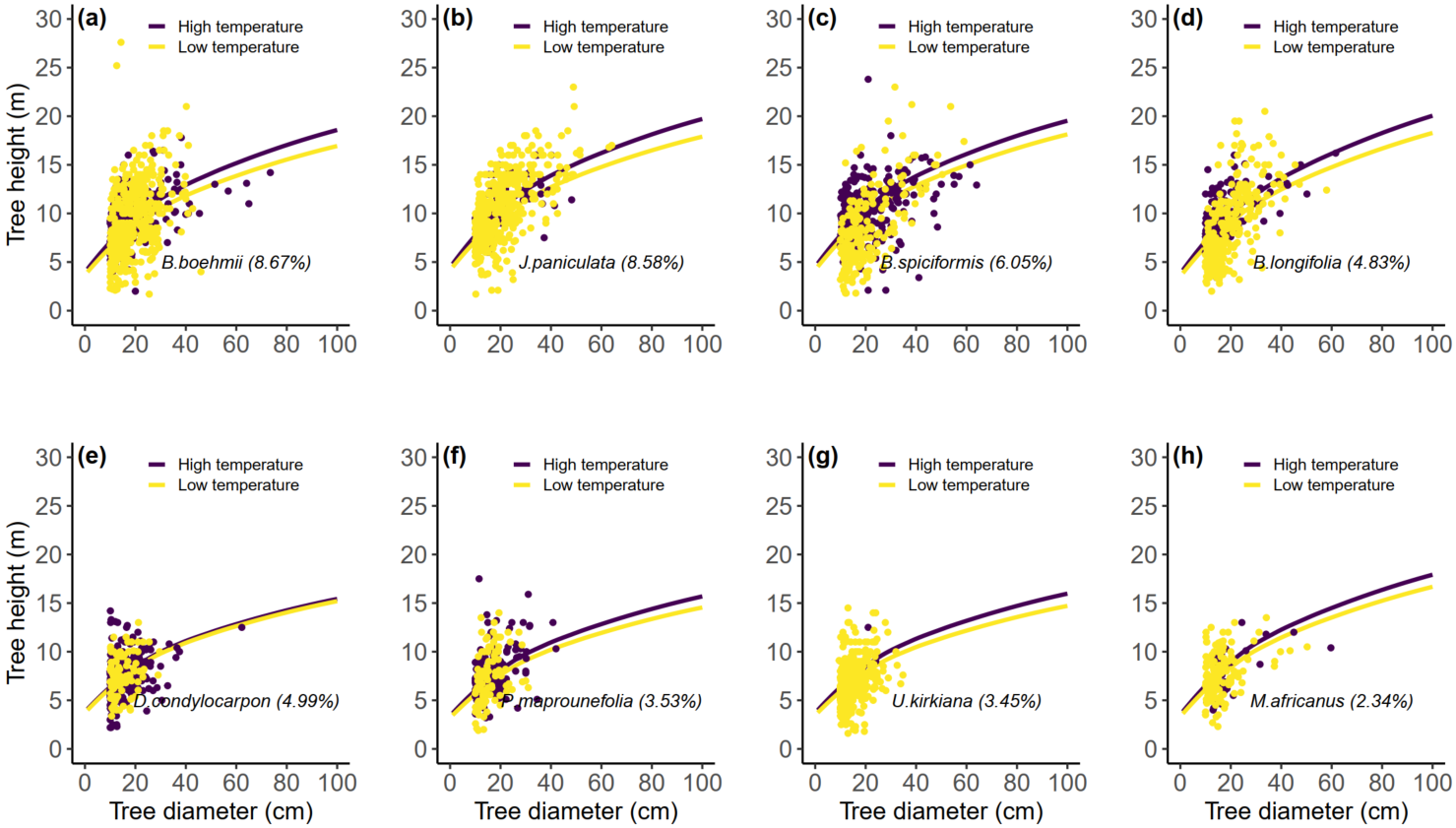

**Fig. S15:** Effects of temperature on intra-specific variability in *H-D* scaling relationships for the four most abundant tall (**a**–**d**) and short (**e**–**h**) stature miombo tree species. Lines represent predictions for one-standard deviation increase (purple) and decrease (yellow) in temperature Circles represent raw data from plots that are either one-standard deviation above (purple) or below (yellow) mean temperature. The percentage in brackets beside each species name corresponds to its relative abundance in the NFI data.

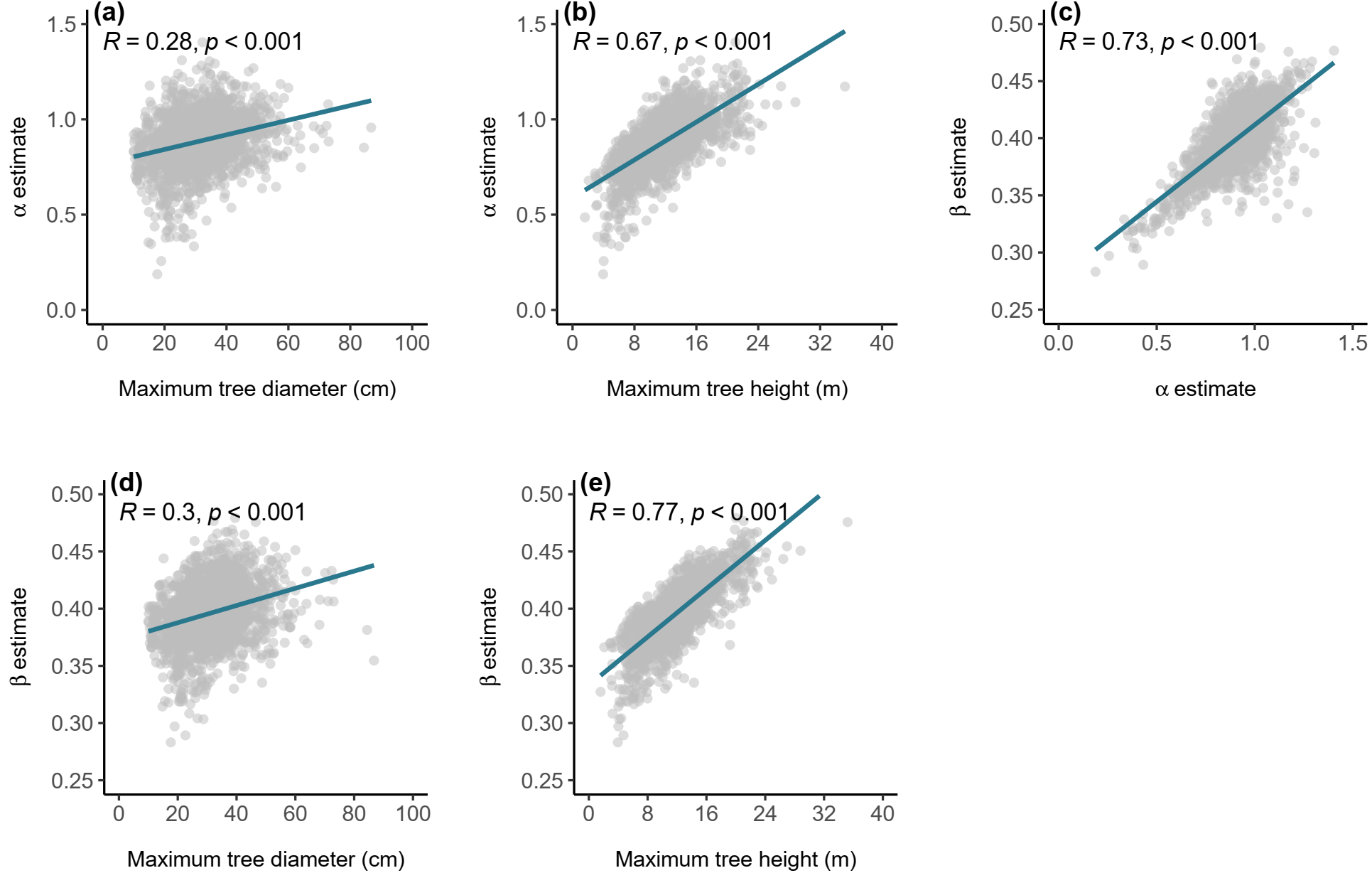

**Fig. S16:** Correlation between estimated (based on equation 3 in the main text) power law model parameters, maximum tree diameter and maximum tree height per plot for the NFI dataset.

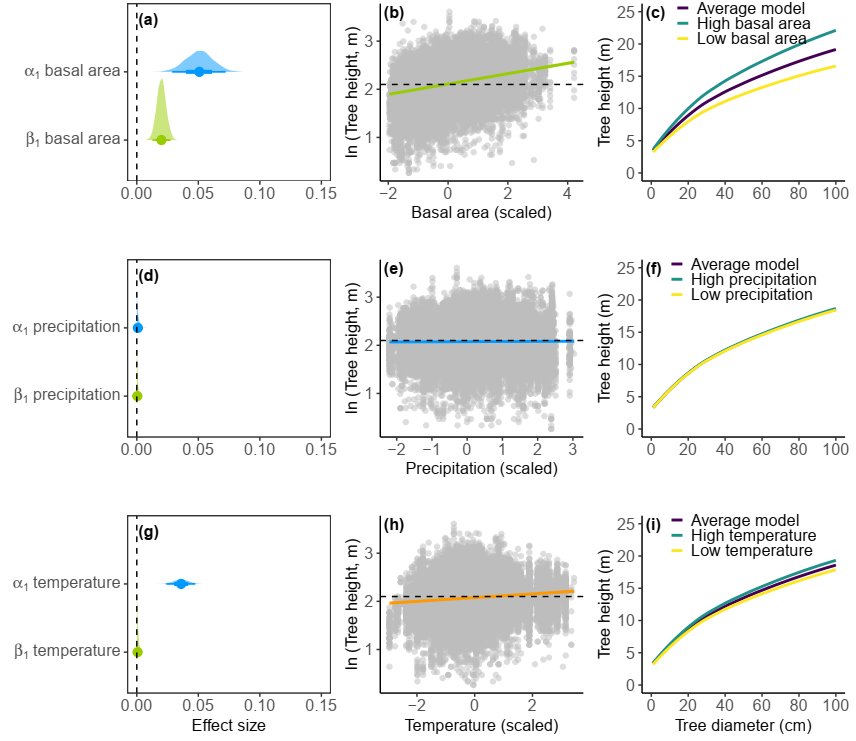

**Fig. S17:** Effects of basal area **(a)** , precipitation **(d)** and temperature **(g)** on the height–diameter (*H-D*) power law model parameters; *H* predictions for a tree of mean size along gradients of basal area **(b)**, precipitation **(e)** and temperature **(h)**; and *H* predictions across a range of *D* values (1-100 cm) for each model predictor, i.e. basal area **(c)**, precipitation **(f)** and temperature **(i)**. In (**b)**, **(e)** and **(h)**, coloured lines represent mean model prediction (with shaded 95% credible intervals), the horizontal dashed line represents the mean tree *H* (on log scale) across the NFI dataset, and data points are shown as grey circles. In (**c)**, **(f)** and **(i)** , low (yellow line) and high (blue line) prediction scenarios correspond to ± 1 standard deviation.

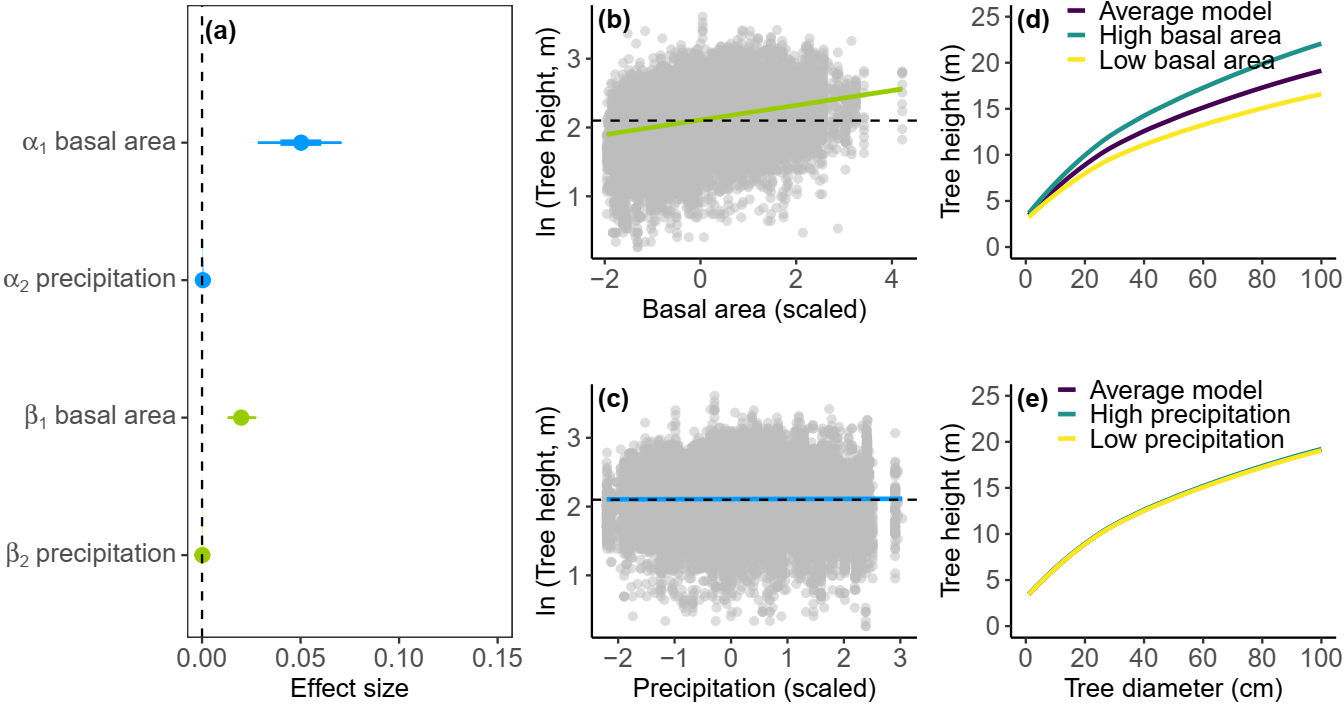

**Fig. S18:** Effects of basal area and precipitation on **(a)** the height–diameter (*H-D*) power law model parameters, **(b**–**c)** *H* predictions for a tree of mean size along gradients of basal area and precipitation, and (**d**–**e**) *H* predictions across a range of *D* values (1-100 cm) while varying one model predictor (e.g., basal area in **d**) and keeping the other one (precipitation) constant at its mean value and setting its effect to zero. In (**b**–**c**), coloured lines represent mean model predictions (with shaded 95% credible intervals), the horizontal dashed line represents the mean tree *H* (on log scale) across the NFI dataset, and data points are shown as grey circles. In (**d**–**e**) low (yellow line) and high (blue line) prediction scenarios correspond to ± 1 standard deviation.

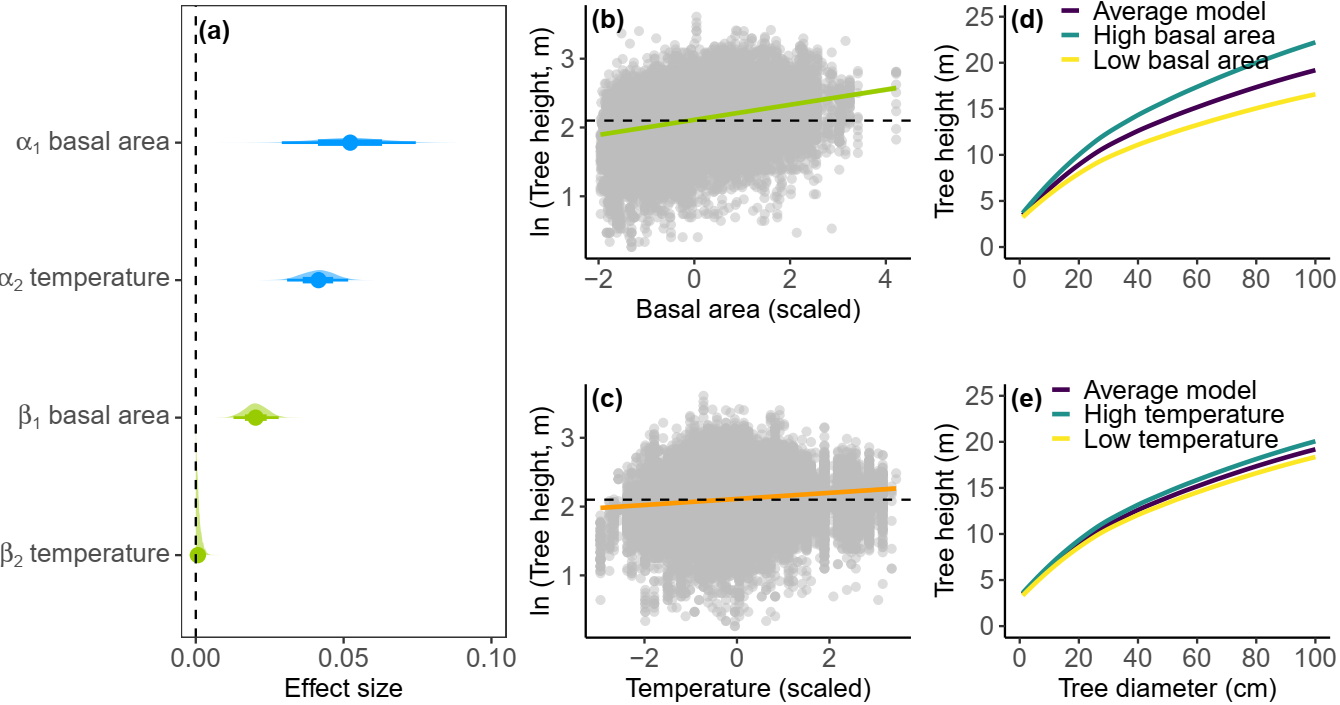

**Fig. S19:** Effects of basal area and temperature on **(a)** the height–diameter (*H-D*) power law model parameters, **(b**–**c)** *H* predictions for a tree of mean size along gradients of basal area and temperature, and (**d**–**e**) *H* predictions across a range of *D* values (1-100 cm) while varying one model predictor (e.g., basal area in **d**) and keeping the other one (temperature) constant at its mean value and setting its effect to zero. In (**b**–**c**), coloured lines represent mean model predictions (with shaded 95% credible intervals), the horizontal dashed line represents the mean tree *H* (on log scale) across the NFI dataset, and data points are shown as grey circles. In (**d**–**e**) low (yellow line) and high (blue line) prediction scenarios correspond to ± 1 standard deviation.

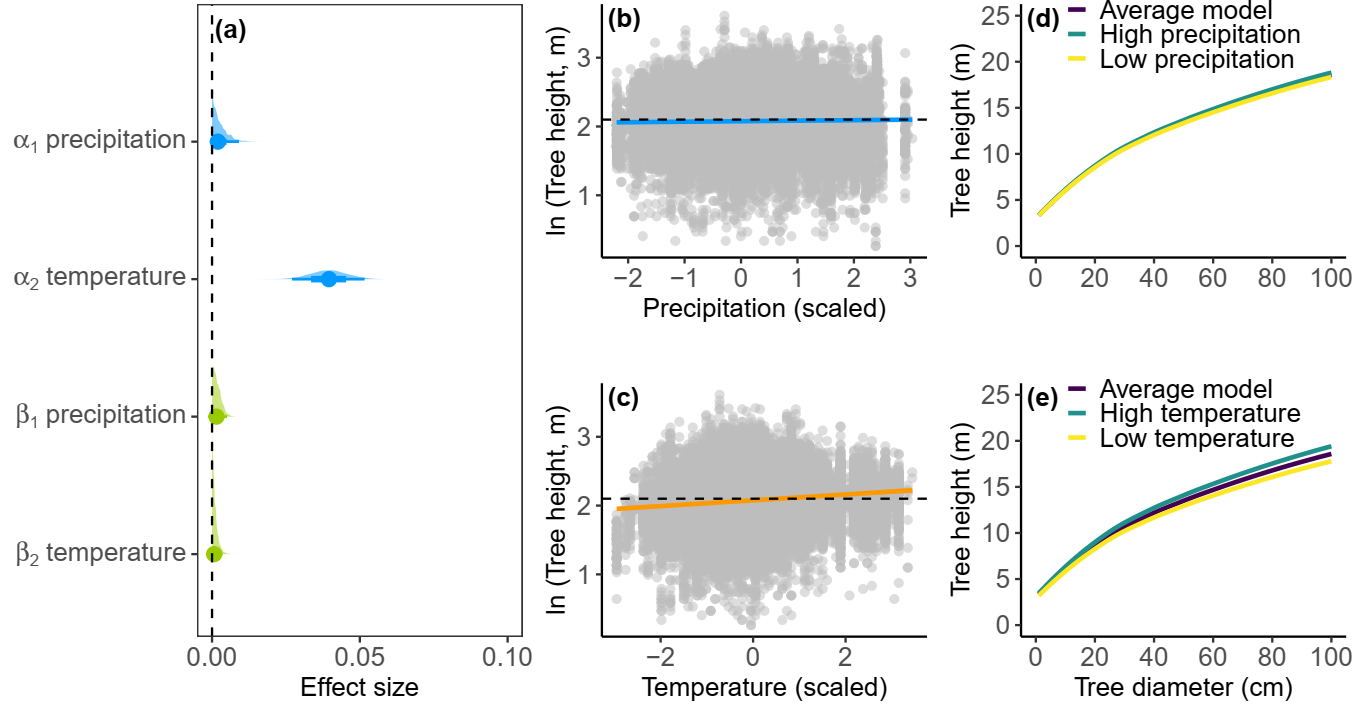

**Fig. S20:** Effects of precipitation and temperature on **(a)** the height–diameter (*H-D*) power law model parameters, **(b**–**c)** *H* predictions for a tree of mean size along gradients of precipitation and temperature, and (**d**–**e**) *H* predictions across a range of *D* values (1-100 cm) while varying one model predictor (e.g., precipitation in **d**) and keeping the other one (temperature) constant at its mean value and setting its effect to zero. In (**b**–**c**), coloured lines represent mean model predictions (with shaded 95% credible intervals), the horizontal dashed line represents the mean tree *H* (on log scale) across the NFI dataset, and data points are shown as grey circles. In (**d**–**e**) low (yellow line) and high (blue line) prediction scenarios correspond to ± 1 standard deviation.

R Core Team ( 2023). _R: A Language and Environment for Statistical Computing_. R

Foundation for Statistical Computing, Vienna, Austria. https://www.R-project.org/.

Schloerke, B., D. Cook, J. Larmarange, F. Briatte, M. Marbach, E. Thoen, A. Elberg and J. Crowley (2021). GGally: Extension to 'ggplot2'. R package version 2.1.2, https://CRAN.R-project.org/package=GGally.

Tredennick, A. T., Bentley, L. P., & Hanan, N. P. (2013). Allometric convergence in savanna trees and implications for the use of plant scaling models in variable ecosystems. *PloS one*, *8*(3), e58241.

Zhou, Y., Bomfim, B., Bond, W. J., Boutton, T. W., Case, M. F., Coetsee, C., Davies, A. B., February, E. C., Gray, E. F., & Silva, L. C. (2023). Soil carbon in tropical savannas mostly derived from grasses. *Nature Geoscience*, *16*(8), 710-716.
